# Human claustrum neurons encode uncertainty and prediction errors during aversive learning

**DOI:** 10.64898/2026.03.13.711590

**Authors:** Mingyue Hu, Rodrigo Dalvit, Mauricio Medina-Pizarro, Maximilian Dougherty, Yajun Zhou, Jonathan Barreto-Nieves, Sami Obaid, Arman Afrasiyabi, Smita Krishnaswamy, Alfred P. Kaye, Murat Gunel, John H. Krystal, Kevin N. Sheth, Xiaosi Gu, Christopher P. Pittenger, Toby Wise, Eyiyemisi C. Damisah

**Author notes:** These authors contributed equally (Mingyue Hu, Rodrigo Dalvit, Mauricio Medina-Pizarro).

## Abstract

Flexible behavior depends on continuous updating of internal models, yet the neural circuits coordinating this process remain poorly understood [1]. The claustrum — reciprocally connected to nearly the entire neocortex — is uniquely positioned to influence cortical processing. Here we report single-neuron recordings from the human claustrum during aversive learning [2], with anterior cingulate cortex and amygdala recordings for comparison. Claustrum and anterior cingulate neurons displayed structured, task-related responses. Distinct subpopulations encoded stimulus onset and action-contingent outcomes, with outcome representations diverging between regions. Critically, both regions encoded model-derived latent variables — uncertainty and prediction error — but with different temporal profiles: only the anterior cingulate carried uncertainty signals during the intertrial period, while both regions encoded uncertainty and prediction error during the active-avoidance period. The amygdala, by contrast, showed minimal latent-variable modulation. These findings provide evidence that human claustrum neurons track higher-order cognitive variables not directly observable from sensory input, and reveal dissociable roles for the claustrum and anterior cingulate cortex in tracking latent task states.

## 1 Introduction

The human brain’s ability to learn and adapt in ever-changing environments emerges from its hierarchical organization and the evolved interplay of distributed neural systems. The claustrum (CLA), a thin sheet-like structure buried deep within subcortical white matter [3], maintains widespread cortical and subcortical connections [4, 5], yet its functional role remains poorly understood. Theoretical accounts of CLA function range from assigning it a central role in conscious integration to dismissing it as an evolutionary vestige [6, 7]. What has been lacking, particularly in humans, is direct neural evidence linking CLA activity to higher-level cognition.

Anatomical tracing in mice and non-human primates demonstrates that the CLA is densely interconnected with frontal cortical regions [3–6, 8]. Notably, cortical areas that communicate via direct cortico-cortical pathways also receive convergent input from bifurcating CLA neurons [3], suggesting that the CLA is positioned to coordinate distributed cortical activity according to contextual and situational demands. Consistent with this circuit architecture, the anterior cingulate cortex (ACC)—a region implicated in higher-order processes including cognitive control[9] and salience processing in humans[10]—has been shown to exert top-down influence on rodent claustral activity during anticipation and task engagement [11, 12]. Reciprocally, ACC signals can be transiently amplified through CLA projections to parietal association and visual cortices [13], and large-scale recordings further indicate that CLA modulates cortical dynamics in a structured, layer-specific manner [13, 14]. Optogenetic stimulation in rodents has also shown that CLA preferentially propagates top-down signals rather than bottom-up signals [12]. In contrast, single-unit recordings from non-human primates [15] suggest that CLA representations are predominantly unimodal, spatially segregated, and lack audiovisual integration — features consistent with the traditional view of the CLA as a passive sensory relay. This discrepancy may reflect, in part, the scarcity of direct neuronal recordings from primates, and the CLA’s small size, which complicates accurate localization with functional neuroimaging. Furthermore, while CLA activity has been extensively recorded in rodents, evidence linking its function to computationally defined task variables remains scarce [11]. This contrasts with the ACC, whose role in higher-order cognition is well established across species, providing a useful reference point [16–20].

To address this gap, we recorded single-unit activity from the CLA, ACC, and amygdala (AMY) in humans performing an aversive learning task [2]. Framed as a visuomotor computer game, the task required participants to steer a spaceship to avoid asteroids while implicitly monitoring stimulus-outcome contingencies. This design permitted approximate Bayesian modeling to derive trial-by-trial estimates of subjective uncertainty and prediction error. By linking these model-inferred variables to neuronal activity, we examined whether and how abstract, task-relevant states are represented in CLA neurons, with ACC and AMY as comparisons.

Our findings reveal that CLA neurons exhibit sharp, time-locked responses to visual stimulus onset and encode prediction error and subjective uncertainty, similar to ACC neurons. However, CLA and ACC differed in the representation of action-contingent outcomes, suggesting region-specific roles in signaling task-relevant information. These findings provide cellular-level evidence in humans that CLA tracks latent task states, implicating it as part of the brain’s inferential hierarchy as envisioned by predictive processing frameworks[1, 21–24]. This engagement in inferential processes may reflect CLA’s distinctive anatomical position, enabling rapid coordination of distributed cortical networks to support timely adaptation to dynamic environmental demands.

## 2 Results

### 2.1 Neural recording and behavioral task

Seven patients with refractory epilepsy underwent robotic implantation of depth electrodes for seizure-onset localization. Each macro-electrode contained eight 40-*µ*m microwires protruding 4mm from the tip, targeting the claustrum (CLA; 4 subjects, 6 microwire bundles), dorsal anterior cingulate cortex (dACC; 6 subjects, 8 bundles), and basolateral amygdala (AMY; 3 subjects, 4 bundles). Precise localization was confirmed through co-registration of preoperative MRI with high-resolution postoperative CT (Fig. 1a-b; Extended Data Figure 1a,b; Extended Data Figure 2a,b; Tables S1 and S2).

**Fig. 1.**
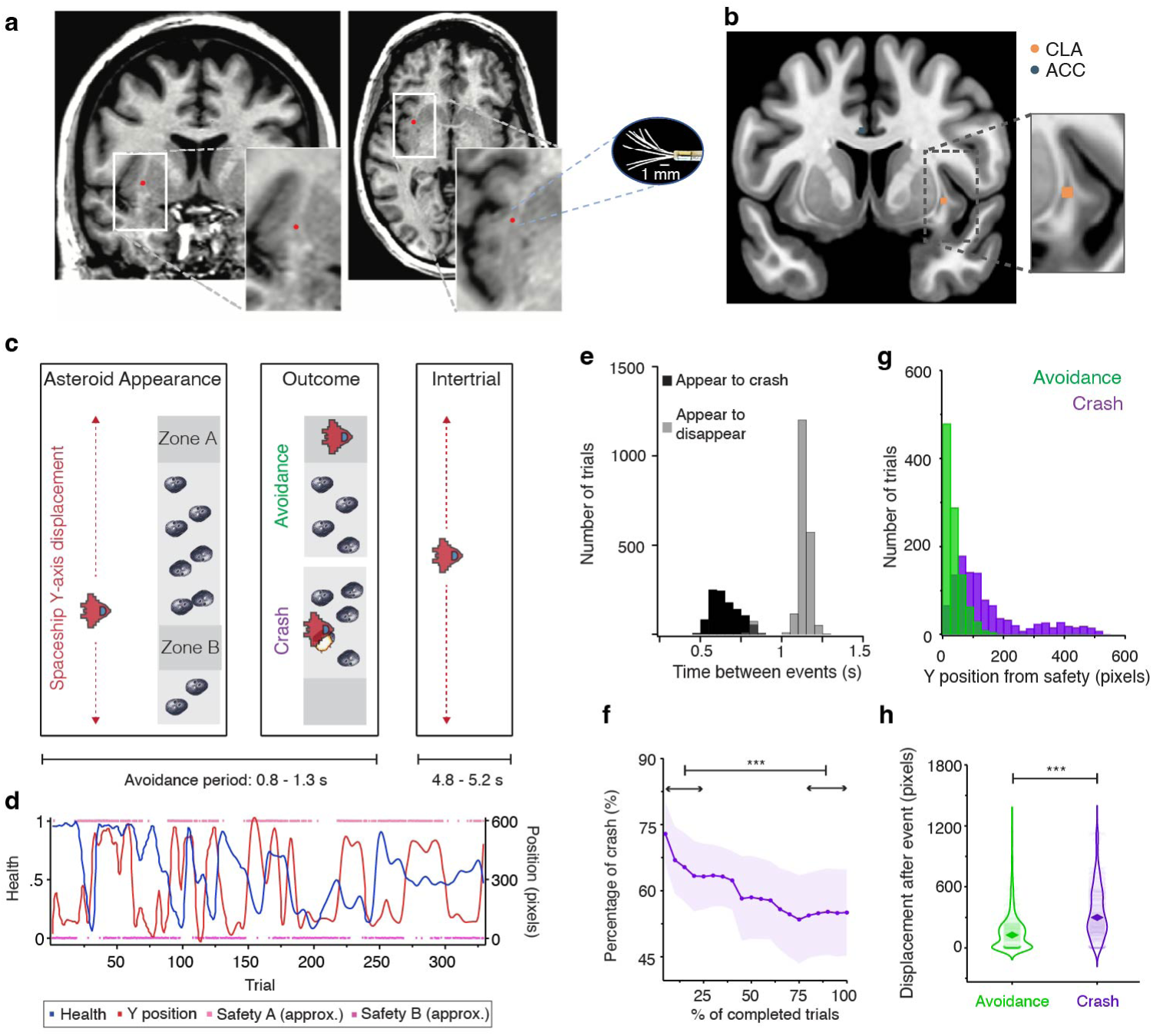
Recording Locations and Task Analysis. (a) MRI brain scans (coronal and axial) co-registered with postoperative CT showing microelectrode locations in the claustrum (CLA). Red dots indicate microwire positions within the 1-mm-thick CLA. Insets show magnified views of the anterior and middle CLA. The oval depicts a magnified view of the microelectrodes used to record neurons. (b) Coronal MNI-152 brain template depicting an example microelectrode location in the CLA (orange) and anterior cingulate cortex (ACC, blue). The inset shows a magnified view of the CLA recording sites. (c) Task structure overview: subjects played a computer-based game involving navigation of a spaceship while avoiding collision with an asteroid belt. Each asteroid belt had two possible safety-zone locations (top and bottom), with at least one location always containing a safety hole, and subjects were instructed to fly through the safety hole to maximize points. Adjustment of the spaceship position was only possible along the *y*-axis, while asteroids moved toward the spaceship on each trial. Successful avoidance increased points (avoidance, green), whereas crashes reduced spaceship integrity (crash, purple). Black horizontal lines illustrate the epochs used for analysis. Boxes indicate the task events analyzed: asteroid appearance, outcome (crash or avoidance), and intertrial. (d) Responses for an example subject across 324 trials. The red line indicates spaceship *y*-position, the blue line indicates spaceship integrity(health), and pink and magenta dots indicate approximate safety zone locations. (e) Distribution of time intervals between asteroid appearance and crash (black), and between asteroid appearance and disappearance (grey). (f) Percentage of crashes as a function of task completion (number of trials) across subjects (1st quarter vs. 4th quarter: *Z* = 4.34, *p* = 1.42 *×* 10^−5^, *n* = 7 subjects, 9 sessions; Wilcoxon signed-rank test). (g) Distribution of *y*-position relative to the closest safety zone at the time of asteroid appearance for crash (purple) and avoidance (green) outcomes across all trials. (h) *y*-position displacement during the first 3 s following crash or avoidance across all subjects (avoidance vs. crash: *Z* = 19.8, *p* = 1.75 *×* 10^−87^; Wilcoxon signed-rank test). *** *p <* 0.001.

Participants controlled a spaceship navigating through approaching asteroids. Two zones (upper, A; lower, B) each had a dynamically changing probability of containing a safe passage. These probabilities were held as high or low (90% vs. 10%) over blocks of 20 to 80 trials before switching, creating a non-stationary environment. Within each zone, the safety hole occupied one of three preset vertical positions, following a pseudo-random sequence fixed across participants and runs. At least one zone was safe on every trial. Because the asteroids approached too quickly for reactive adjustment, successful avoidance required anticipatory positioning based on learned probabilistic beliefs. Failure resulted in a crash and a loss of spaceship health (integrity) (Fig. 1c,d). This structure enabled computational inference of participants’ internal beliefs about safety under uncertainty, defined as the estimated probability that a safety hole would be present within each zone. Beliefs about the safety probability of each location were modeled using an approximate Bayesian framework, in which beta distributions were updated trial-by-trial based on the observed outcome (presence or absence of a safety hole). Uncertainty was derived from the posterior variance of the beta distribution and prediction error from the mismatch between observed outcomes and estimated safety probability.

Participants completed 227 trials on average (range: 125–324). Each trial lasted 6s. Within each trial, the active avoidance (i.e. asteroids visible on the screen) epoch lasted 1.1 s on average (range: 0.8–1.3 s), and the interval from asteroid onset to the action-contingent outcome (successful avoidance or crash) averaged 0.65 s (range: 0.48–0.9 s). The remaining trial time comprised the intertrial interval, during which no asteroids were present on the screen, lasting 5 s on average (range: 4.8–5.2 s). This relatively brief interval constrained deliberation and encouraged participants to engage in anticipatory spaceship positioning (Fig. 1e).

All subjects engaged robustly with the task. Performance improved with practice, as crash rates dropped from 66 *±* 2.9% to 54.9 *±* 3.6% between the first 25% and the last 25% of trials (*p* = 1.42 *×* 10*^−^*^5^, Wilcoxon signed-rank; Fig. 1f). Crashes triggered behavioral adjustments—subjects repositioned 2.2× more after a crash than after asteroid avoidance (364.2 *±* 8 vs 162.7 *±* 6 pixels [mean *±* s.e.m], *p* = 1.75 *×* 10*^−^*^87^, Wilcoxon signed-rank test, Fig. 1h), suggesting rapid strategy updates following aversive action-contingent outcome.

We next examined neuronal responses across the corresponding trial epochs in the CLA and ACC. Overall, the proportion of task-responsive neurons increased sharply after asteroid onset in both regions, but peaked earlier in CLA than in ACC, indicating earlier recruitment of CLA populations following stimulus onset (Fig. 2c,d).

**Fig. 2.**
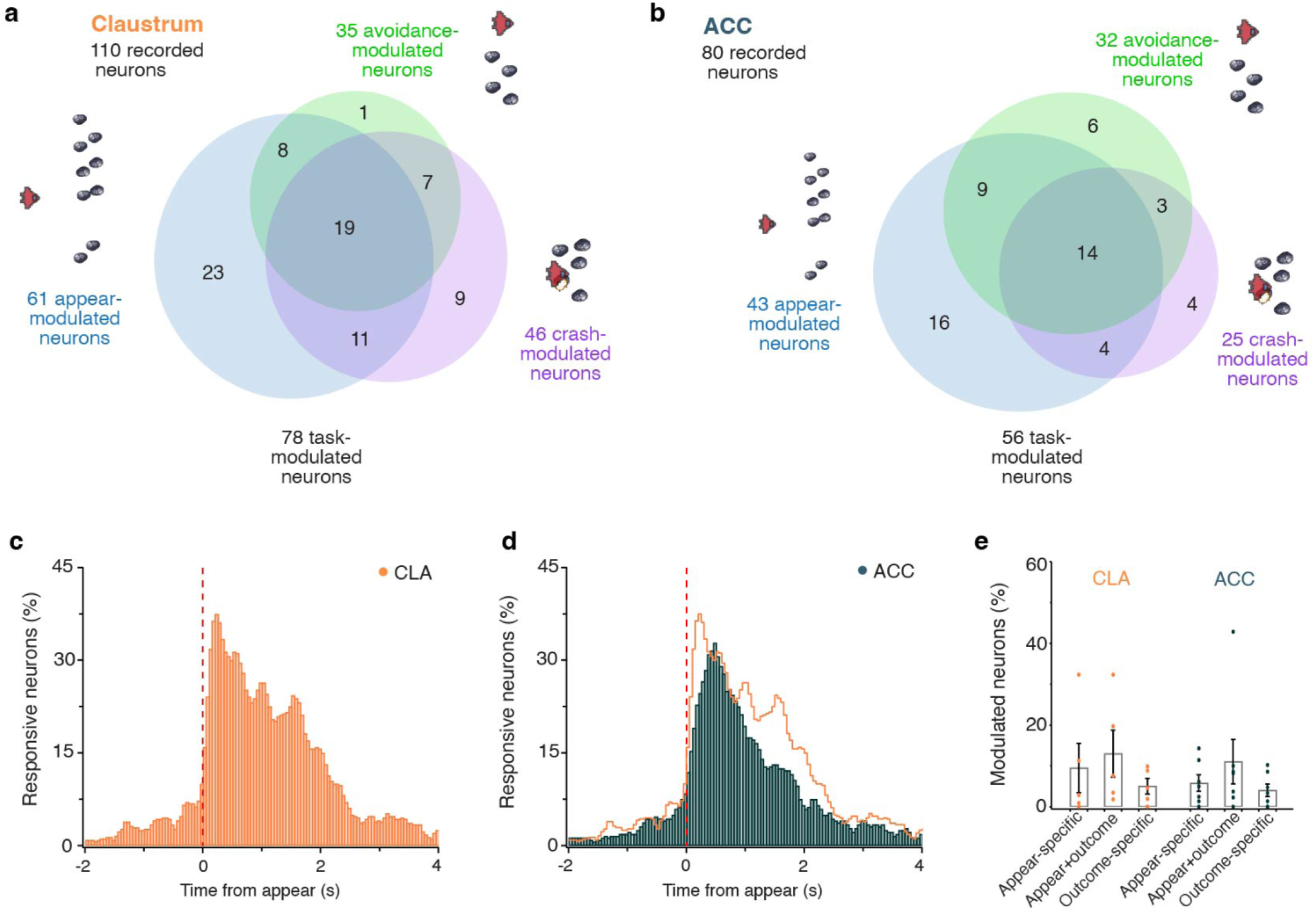
Distribution and timing of CLA and ACC responsive neurons. (a) Venn diagram illustrating the distribution of claustrum (CLA) neurons, showing task-responsive units (78/110) across the three analyzed task events: asteroid appearance (blue), avoidance (green), and crash (purple). (b) Same as (a), but for anterior cingulate cortex (ACC) neurons, showing the distribution of task-modulated single units (56/80). (c) Fraction of responding CLA neurons (percentage of total recorded units) as a function of time relative to asteroid appearance. (d) Same as (c), but for ACC neurons. Note that the peak fraction of responding neurons occurs earlier for CLA than for ACC. (e) Percentage of responding neurons per recording session (dots) with corresponding medians (bars) for each classified neuron type, shown separately for CLA (orange) and ACC (blue). Recording session-level percentages did not differ significantly across neuron types within either region (CLA, *n* = 5 sessions, *χ*^2^ = 0.85, *p* = 0.65, two-sided Kruskal-Wallis test; ACC, *n*= 7 sessions, *χ*^2^ = 1.07, *p* = 0.59, two-sided Kruskal-Wallis test).

### 2.2 CLA neurons show diverse task-related firing patterns

78 of the 110 recorded CLA single neurons (71%) showed significant firing changes during task events (Fig. 2a, 3d). Among these, 61 neurons showed early firing-rate changes following asteroid appearance (Fig. 2a, 3a,b,e) comprising neurons that increased (*n* = 39, bursters) or decreased (*n* = 22, pausers) their firing rate compared to baseline.

**Appear-specific responses:** 21% of recorded units (Fig. 2a) showed predominantly increased firing after asteroid appearance (Δfiring rate, pre-appearance: 0.11 *±* 0.02 vs post-appearance: 0.62 *±* 0.13 spikes/s [mean *±* s.e.m], *p* = 2.7 *×* 10*^−^*^5^, Wilcoxon signed-rank, *n* = 23; Fig. 3f; Fig. 3g left) without significant firing rate changes during action-contingent outcome (avoidance: 0.12 *±* 0.02 vs crash: 0.18 *±* 0.04 spikes/s [mean *±* s.e.m], *p* = 0.14, Wilcoxon signed-rank test, *n* = 23; Fig. 3g right). Appear-specific neurons responded selectively to asteroid appearance but not to other visual stimuli such as spaceship displacement (Extended Data Figure 4a), indicating that they were not broadly responsive to visual motion or positional updates. We therefore speculate that this response pattern may reflect a sensitivity to behaviorally salient or unpredictable events, as previously reported in non-human primates [15].

**Fig. 3.**
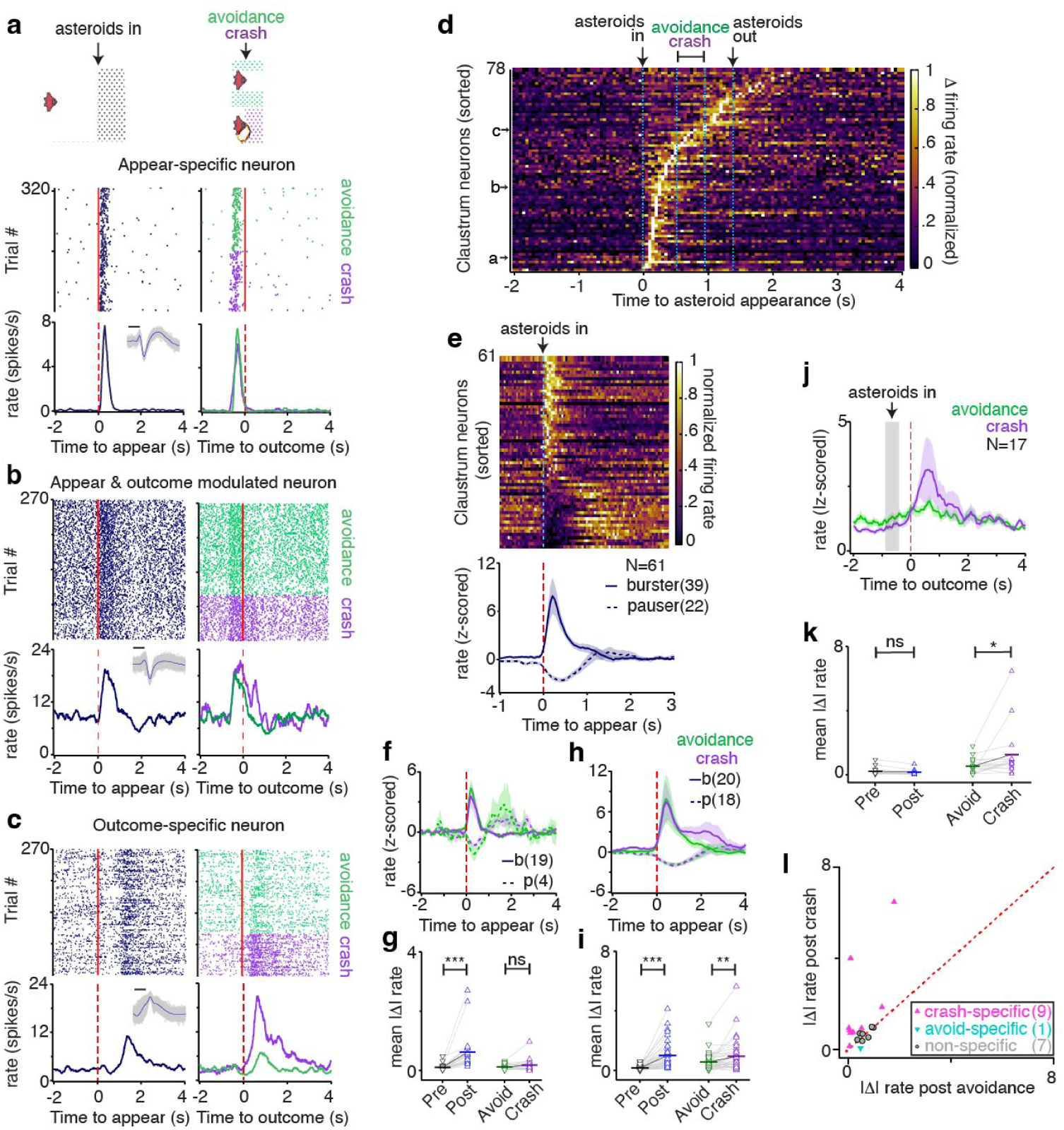
Claustrum neuronal response to asteroid appearance and outcome. (a) Top: task schematic illustrating trial segmentation based on asteroid appearance and action-contingent outcome (crash or avoidance). Middle and bottom: raster plot and firing rate of an example appearance-specific neuron. (b) Raster plot and firing rate of an example appearance– and outcome-modulated neuron in the claustrum (CLA). (c) Raster plot and firing rate of an example outcome-modulated neuron in the CLA. (d) Heatmap showing normalized firing rate changes for all task-modulated CLA neurons aligned to asteroid appearance. Example neurons shown in panels a–c are indicated by the corresponding labels on the y-axis. (e) Top: heatmap of normalized firing rates for all appearance-modulated CLA neurons with bursters shown on top and pausers on the bottom (*n* = 61). Bottom: average responses (z-score *±* s.e.m.) for appearance-modulated CLA neurons, categorized as bursters (*n* = 39, solid line) and pausers (*n* = 22, dashed line), aligned to asteroid appearance. (f) Average responses (z-score *±* s.e.m.) for appearance-specific CLA neurons, categorized as bursters (solid lines) and pausers (dashed lines). (g) Mean absolute change in firing rate for appearance-specific CLA neurons. Left: pre– vs. post-appearance (*n* = 23, *Z* = *−*4.19, *p* = 2.7 *×* 10^−5^). Right: post-crash vs. post-avoidance (*n* = 23, *Z* = 1.46, *p* = 0.14). (h) Average responses (z-score *±* s.e.m.) for appearance-and outcome-modulated CLA neurons separated by outcome (avoidance, green; crash, purple) (i) Mean absolute change in firing rate for appearance– and outcome-modulated CLA neurons. Left: pre-vs. post-appearance (*n* = 38, *Z* = *−*5.37, *p* = 7.74 *×* 10^−8^). Right: post-crash vs. post-avoidance (*n* = 38, *Z* = 2.83, *p* = 0.0046). (j) Average responses (z-score *±* s.e.m.) for all outcome-modulated CLA neurons (*n* = 17) aligned to crash or avoidance. (k) Mean absolute change in firing rate for outcome-specific CLA neurons. Left: pre– vs. post-appearance (*n* = 17, *Z* = 1.21, *p* = 0.23). Right: post-crash vs. post-avoidance (*n* = 17, *Z* = 2.53, *p* = 0.01). (l) Scatter plot of mean absolute firing rate change post-avoidance (x-axis) versus post-crash (y-axis) for outcome-specific CLA neurons (pink triangles; *n* = 9), avoidance-specific (blue triangle; *n* = 1), and non-specific (grey dots; *n* = 7). Two-sided Wilcoxon signed-rank tests were used to assess firing rate changes, with significance levels indicated for FDR-corrected *p* values (*** *p <* 0.001, ** *p <* 0.01, * *p <* 0.05, n.s., not significant).

**Appear and outcome responses:** Thirty-eight neurons (35% of units) exhibited biphasic firing: gradual increases after asteroid appearance and secondary elevations during action-contingent outcomes. These “appear-and-outcome” neurons showed significant increases after appearance (0.17 *±* 0.03 vs 1.0 *±* 0.15 spikes/s [mean *±* s.e.m], *p* = 7.74*×*10*^−^*^8^, Wilcoxon signed-rank test, *n* = 38; Fig. 3b,h,i) and stronger responses to crashes than avoidance (crash: 0.94 *±* 0.17 vs avoidance: 0.58 *±* 0.09 spikes/s [mean *±* s.e.m], *p* = 0.0046, Wilcoxon signed-rank test, *n* = 38; Fig. 3i right). This sustained profile suggests integration of task-relevant information across trial phases.

**Outcome-specific responses:** Seventeen neurons (15% of units) responded exclusively to action-contingent outcomes. These neurons showed no changes at asteroid appearance (0.22 *±* 0.06 vs 0.17 *±* 0.04 spikes/s [mean *±* s.e.m], *p* = 0.23, Wilcoxon signed-rank test, *n* = 17; Fig. 3c,j, and k left) but showed stronger activity following crashes than avoidance at the population level (1.27 *±* 0.39 vs 0.54 *±* 0.12 spikes/s [mean *±* s.e.m], *p* = 0.01, Wilcoxon signed-rank test, *n* = 17; Fig. 3k right). This population comprised crash-specific (*n* = 9), avoidance-specific (*n* = 1), and non-specific units (*n* = 7; Fig. 3l), with a bias toward crash-related responses, suggesting specialized outcome-related processing circuits that may support context-dependent monitoring in the CLA.

### 2.3 ACC neurons show task-phase selectivity

The ACC is hypothesized to encode and update internal task-state representations that are necessary for guiding adaptive behavior[16, 17, 25]. We therefore examined neuronal responses in the ACC using the same analytical approach. 56 of the 80 recorded units (70%) were task-responsive (Fig. 2b). Like CLA, ACC neurons increased their firing rate during different phases of each trial (Extended Data Figure 5c).

**Appear-specific responses:** The ACC had neurons that were modulated by asteroid appearance, including bursters (*n* = 30) and pausers (*n* = 13) (Fig. 4d). 16 appear-specific neurons (20% of units) responded selectively to asteroid appearance (0.19 *±* 0.04 vs 0.83 *±* 0.18 spikes/s [mean *±* s.e.m], *p* = 4.38 *×* 10*^−^*^4^, Wilcoxon signed-rank test, *n* = 16; Fig. 4e, 4f left) but did not display any outcome preference (avoidance: 0.32 *±* 0.08 vs crash: 0.22 *±* 0.03 spikes/s [mean *±* s.e.m], *p* = 0.33, Wilcoxon signed-rank test, *n* = 16; Fig. 4f right).

**Fig. 4.**
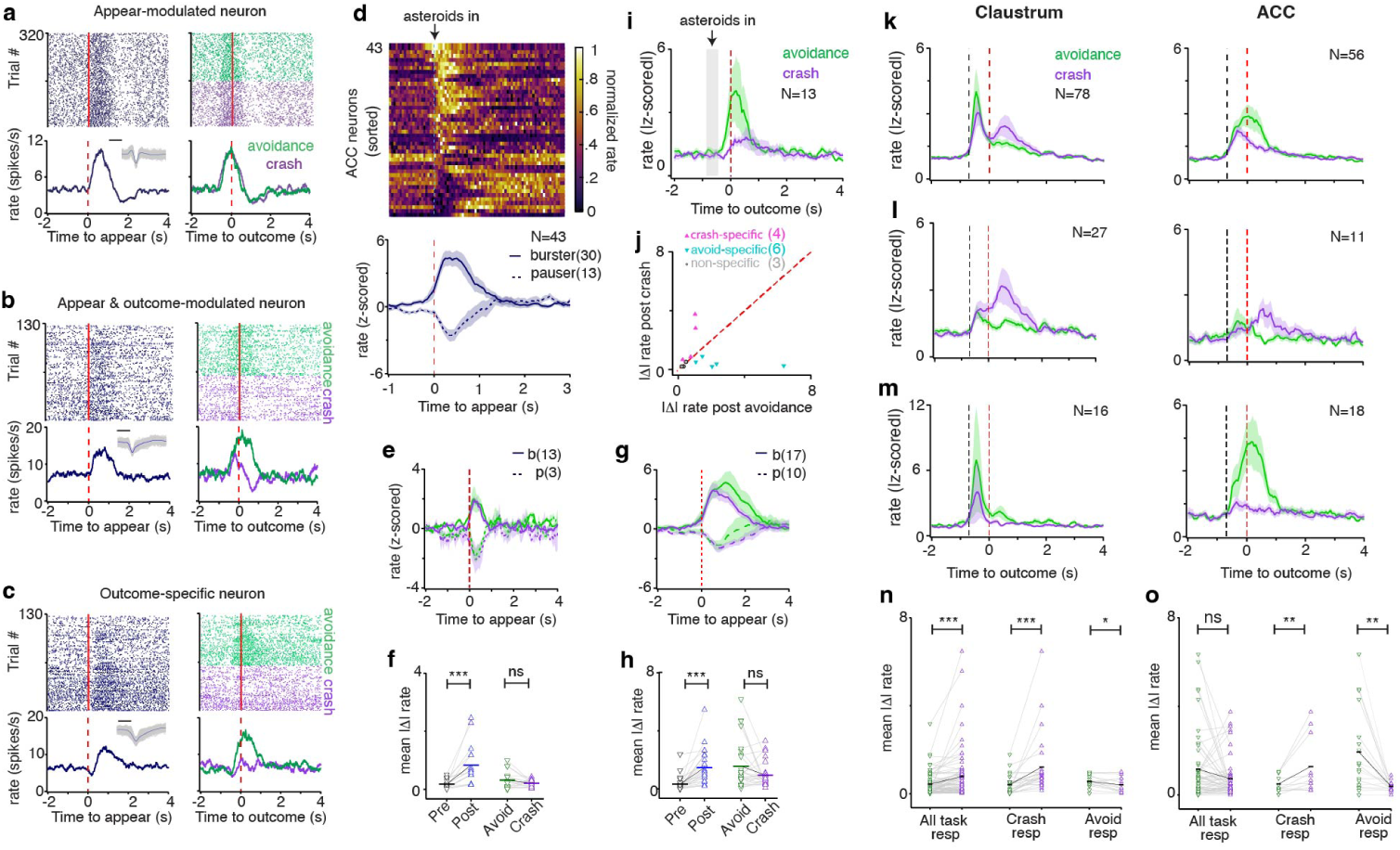
ACC neuronal responses to task and differential CLA and ACC response patterns. (a) Example appearance-specific anterior cingulate cortex (ACC) neuron without action-contingent outcome differentiation. (b) Example appearance– and outcome-modulated ACC neuron. (c) Example outcome-modulated ACC neuron showing increased firing during avoidance (green) but not crashes (purple). (d) Heatmap of normalized firing rates for appearance-modulated ACC neurons (*n* = 43), with bursters shown on top and pausers on the bottom. Bottom panel shows average responses (*±* s.e.m.) for appearance-modulated neurons, categorized as bursters (*n* = 30, solid line) and pausers (*n* = 13, dashed line). (e) Average responses (*±* s.e.m.) for appearance-specific ACC neurons. (f) Mean absolute change in firing rate for appearance-specific ACC neurons. Left: pre– vs. post-appearance (*n* = 16, *Z* = *−*3.52, *p* = 4.38 *×* 10^−4^). Right: post-crash vs. post-avoidance (*n* = 16, *Z* = *−*0.98, *p* = 0.33). (g) Average responses (*±* s.e.m.) for appearance– and outcome-modulated ACC neurons. (h) Mean absolute change in firing rate for appearance– and outcome-modulated ACC neurons. Left: pre– vs. post-appearance (*n* = 27, *Z* = *−*4.52, *p* = 6.28 *×* 10^−6^). Right: post-crash vs. post-avoidance (*n* = 27, *Z* = *−*0.6, *p* = 0.55). (i) Average responses (*±* s.e.m.) for outcome-specific ACC neurons (*n* = 13) aligned to crash or avoidance. (j) Scatter plot of mean absolute firing rate change post-avoidance (x-axis) versus post-crash (y-axis), classifying outcome-specific ACC neurons as crash-specific (pink; *n* = 4), avoidance-specific (blue; *n* = 6), and non-specific (grey; *n* = 3). (k) Average responses (*±* s.e.m.) of task-modulated claustrum (CLA) neurons (left, *n* = 78) and ACC neurons (right, *n* = 56) aligned to outcome. (l) Average responses (*±* s.e.m.) of crash-modulated CLA (left, *n* = 27) and ACC neurons (right, *n* = 11). (m) Average responses (*±* s.e.m.) of avoidance-modulated CLA (left, *n* = 16) and ACC neurons (right, *n* = 18). (n) Mean absolute change in firing rate for CLA neurons in response to avoidance (green) versus crash (purple). Left: all task-responsive neurons (*n* = 78, *Z* = 3.74, *p* = 1.85 *×* 10^−4^). Middle: crash-responsive neurons (*n* = 27, *Z* = 4.16, *p* = 3.23 *×* 10^−5^). Right: avoidance-responsive neurons (*n* = 16, *Z* = *−*2.07, *p* = 0.039). (o) Mean absolute change in firing rate for ACC neurons in response to avoidance (green) versus crash (purple). Left: all task-modulated neurons (*n* = 56, *Z* = *−*1.14, *p* = 0.25). Middle: crash-modulated neurons (*n* = 11, *Z* = 2.67, *p* = 0.005). Right: avoidance-modulated neurons (*n* = 18, *Z* = *−*3.30, *p* = 0.001). Two-sided Wilcoxon signed-rank tests were used, with significance levels indicated for FDR-corrected *p* values (*** *p <* 0.001, ** *p <* 0.01, * *p <* 0.05, n.s., not significant).

**Appear and outcome responses:** Twenty seven neurons (34% of units) were responsive to both asteroid appearance and outcomes. These neurons responded to asteroid appearance (0.34 *±*0.1 vs 1.47 *±*0.22 spikes/s [mean *±* s.e.m], *p* = 6.28 *×*10*^−^*^6^, Wilcoxon signed-rank test, *n* = 27; Fig. 4g, 4h left), but did not display population-level discrimination between outcomes (avoidance: 1.57 *±* 0.32 vs crash: 0.95 *±* 0.16 spikes/s [mean *±* s.e.m], *p* = 0.55, Wilcoxon signed-rank test, *n* = 27; Fig. 4g, 4h right).

**Outcome specific responses:** Thirteen outcome-specific neurons (16% of units) responded exclusively to action-contingent outcomes, with nominally stronger avoidance than crash responses (avoidance: 1.32 *±* 0.46 vs crash: 0.87 *±* 0.31 spikes/s [mean *±* s.e.m], *p* = 0.57, Wilcoxon signed-rank test, *n* = 13; Fig. 4c,i). This population comprised crash-specific (*n* = 4), avoidance-specific (*n* = 6), and non-specific units (*n* = 3; Fig. 4j). Neither appear-specific nor appear-and-outcome ACC neurons responded to spaceship displacement (Extended Data Figure 4d,e).

These results suggest that ACC neurons exhibited heterogeneous but structured task-related responses. Notably, many ACC neurons responded to asteroid appearance via sustained activity, in contrast to the transient activation observed in CLA units (see Fig. 2 for full breakdown of units). This response profile potentially reflects the maintenance of task-state or salience representations [26], in line with context/strategy mapping views [17]. Action-contingent outcome ACC neurons did not show significant discrimination between crash and avoidance outcomes at the population level. Instead, substantial variability was observed across units (Fig. 4h, right), with some neurons firing strongly after crash and others after asteroid avoidance. This variability is consistent with prior reports that ACC encoding may be distributed across distinct, differently tuned subpopulations rather than reflecting a uniform population preference [27].

### 2.4 Distinct outcome patterns between CLA and ACC

Motivated by the difference in outcome preferences between regions, we examined population responses in all task-responsive neurons. This analysis revealed outcome discrimination in CLA but not ACC (CLA crash: 0.79 *±* 0.13 vs avoidance: 0.44 *±* 0.06 spikes/s [mean *±* s.e.m], *p* = 1.85 *×* 10*^−^*^4^, Wilcoxon signed-rank test, *n* = 78; ACC crash: 0.74 *±* 0.12 vs avoidance: 1.16 *±* 0.2 spikes/s [mean *±* s.e.m], *p* = 0.25, Wilcoxon signed-rank test, *n* = 56; Fig. 4k, 4n left, 4o left).

A linear mixed-effects model with region, outcome, and their interaction as fixed effects, and random intercepts for subject and neuron (task-responsive units), revealed a significant region *×* outcome interaction (*β* = 0.771, 95% CI [0.365, 1.177], SE = 0.207, *z* = 3.724, *p <* 0.001). A likelihood-ratio test confirmed that this interaction term significantly improved model fit (*χ*^2^(1) = 20.55, *p* = 5.82 *×* 10*^−^*^6^). This interaction remained significant in leave-one-subject-out refits (all *p <* 0.005) and after log-transforming firing rates (*β* = 0.297, SE = 0.073, *p* = 4.84 *×* 10*^−^*^5^), indicating that the effect was robust to both subject-level sampling variability and distributional skew. These results support a regional dissociation in outcome-related firing-rate modulation. Post hoc simple-effects contrasts showed that CLA firing rates were significantly lower for avoidance than crash outcomes (*β* = *−*0.345, 95% CI [*−*0.607*, −*0.082], SE = 0.134, *z* = *−*2.574, *p* = 0.010), whereas ACC firing rates were significantly higher for avoidance than crash outcomes (*β* = 0.427, 95% CI [0.117, 0.736], SE = 0.158, *z* = 2.700, *p* = 0.0069).

These results diverge from the non-parametric paired comparison, which detected significant outcome-related discrimination in CLA but not in ACC. To examine the basis of this discrepancy, we quantified neuron-wise outcome preference as Δ = FR_avoidance_ *−* FR_crash_. CLA neurons displayed mixed-sign tuning, but with a clear population level bias (27/78 avoidance-preferring, 51/78 crash-preferring; sign test, *p* = 0.0088). By contrast, ACC neurons were substantially more heterogeneous, with nearly balanced outcome preferences across units (27/56 avoidance-preferring, 28/56 crash-preferring, 1/56 no difference; sign test, *p* = 1.0). Thus, although ACC preferred one outcome or the other, the absence of a consistent directional bias across the population explains why the Wilcoxon test was not significant despite a mean-based mixed-effects model detecting an average outcome effect.

Given these observations, we hypothesized that this heterogeneity may reflect sensitivity to belief-related latent variables such as prediction error and uncertainty, which have previously been linked to ACC activity [18, 20, 25, 27–30]. As our central goal was to determine whether similar latent-variable coding could be identified in the CLA, we first examined this possibility in CLA neurons and then applied the same model-based analysis to ACC.

### 2.5 CLA neurons reflect hidden task variables

We examined whether single neurons contained information about latent states—uncertainty and prediction error (PE)—derived from an approximate Bayesian model (ALB-sticky; see Methods) [2]. We hypothesized that spaceship position changes relative to inferred safety probability (Fig. 1d, 5a) may indicate implicit tracking of latent features. To examine whether model derived uncertainty had a behavioral read-out, we stratified movement trajectories using the top and bottom 30th percentile cutoffs for model-inferred uncertainty (for safety zones A and B separately). Movement magnitude was quantified as the mean per-trial sum of absolute *y*-position changes within –4 to 0 s window aligned to asteroid appearance (low: 118.31 *±* 47.78 vs high: 146.40 *±* 42.87, mean *±* SD; *Z* = *−*2.35, *p* = 0.017; Fig. 5b, left), and movement variability as the mean per-trial standard deviation of *y*-position (low: 23.72 *±* 13.28 vs high: 29.33 *±* 11.92, mean *±* SD; *Z* = *−*2.48, *p* = 0.011; Fig. 5b, right). Under low-uncertainty conditions, participants exhibited less movement and lower within-trial variability. In contrast, high-uncertainty trials were associated with larger and more variable movements, suggesting increased exploratory-related behavior between potential safety zones. These movement differences suggest that model-derived uncertainty may capture behaviorally meaningful variation, supporting its use as a proxy for participants’ internal states.

**Fig. 5.**
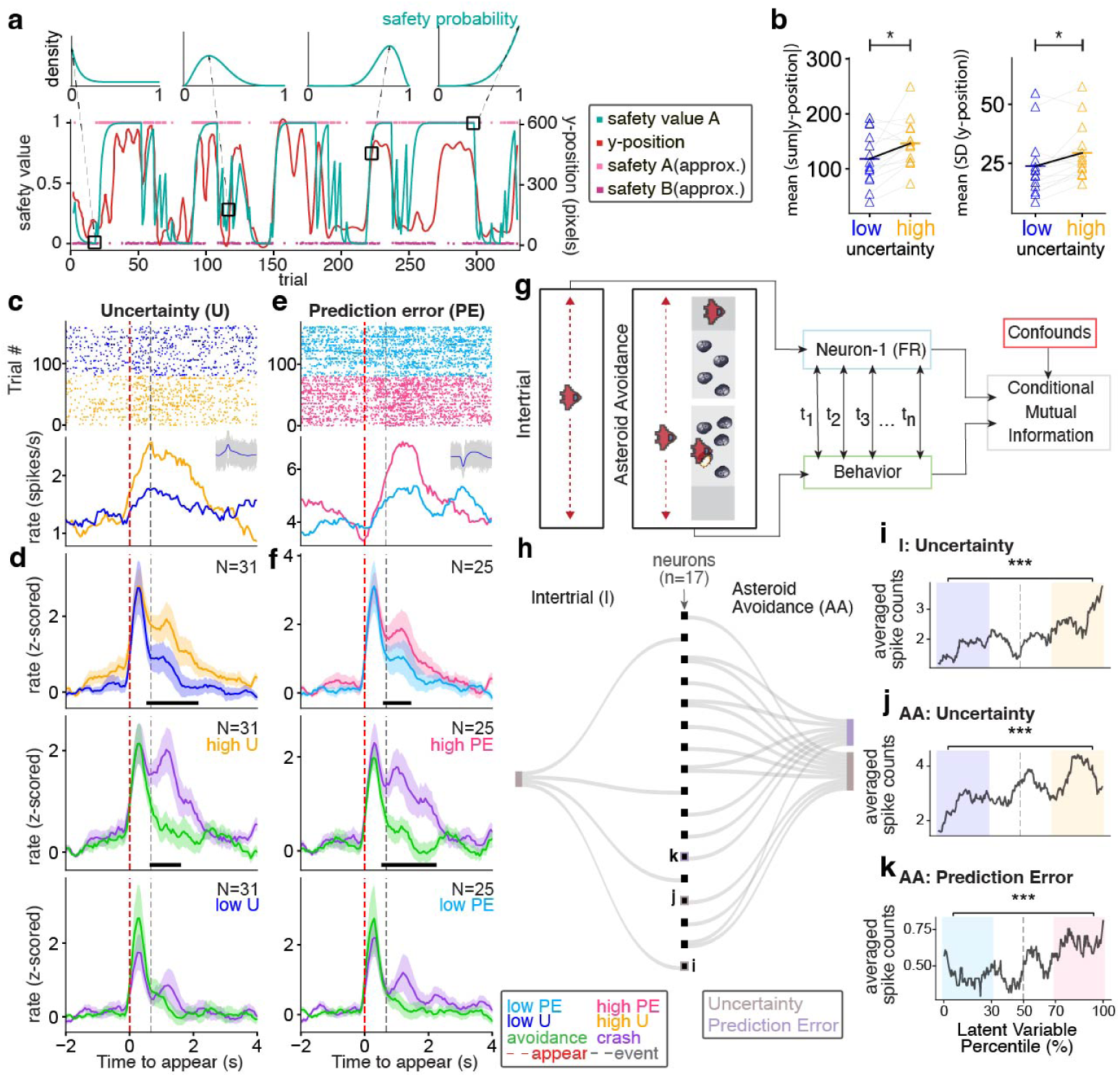
CLA neurons track uncertainty and prediction error. (a) Top: Example beta distributions illustrating trial-wise safety probability and uncertainty (posterior variance). The *x*-axis spans safety probabilities (0–1), and the *y*-axis indicates probability density. Dashed lines and solid boxes denote the distribution mean, used as the model’s safety estimate per trial, guiding behavior and informing prediction error after outcome. Wider distributions reflect greater uncertainty. Bottom: Traces from a representative subject showing estimated safety probability for safety zone A (cyan, safety value), spaceship *y*-position (red), and approximate A/B safety zones (bright/dark pink). Note *y*-position changes following safety value A. (b) Left, mean of the per-trial sum of absolute *y*-position changes (in the 4 s preceding asteroid appearance) under low vs. high uncertainty (low: 118.31 *±* 47.78 vs high: 146.40 *±* 42.87, mean *±* SD; *Z* = *−*2.35, *p* = 0.017, Wilcoxon signed-rank test); Right, mean of the per-trial standard deviation (SD) of *y*-position (low: 23.72 *±* 13.28 vs high: 29.33 *±* 11.92, mean *±* SD; *Z* = *−*2.48, *p* = 0.011, Wilcoxon signed-rank test). Colored triangles represent individual subjects; black lines denote subject-level population means (blue: low uncertainty; orange: high uncertainty). (c, e) Exemplar neurons modulated by uncertainty (c; same color code as in (b)) and prediction error (e; low PE, light blue; high PE, pink). Neurons were identified based on responses during high-uncertainty or high-prediction-error trials. Top: Spike rasters aligned to task events. Bottom: Mean firing rate (*±* s.e.m.) and waveform of the same neuron. (d, f) Population responses of neurons identified as high-uncertainty–modulated (d; *n* = 31) or high-prediction-error–modulated (f; *n* = 25). Top: Z-scored firing rates (*±* s.e.m.) comparing high versus low uncertainty (d) or high versus low prediction error (f) trials. Middle: Responses during high-uncertainty or high-prediction-error trials, with trials separated by action-contingent outcome (avoidance, green; crash, purple). Bottom: Responses during low-uncertainty or low-prediction-error trials, again separated by outcome (avoidance vs crash). (d) Top: *t* = 4.28; middle: *t* = *−*4.89; peak cluster t-statistics (f) top: *t* = 3.08; middle: *t* = *−*3.96 peak cluster t-statistics. Black bars indicate time periods during which neural responses differed significantly between conditions, identified using two-sided cluster-based permutation test (*p <* 0.05). (g) Schematic illustrating conditional mutual information (MI) analysis computed on a trial-wise basis between latent variable and firing rate, during the intertrial (left) and asteroid avoidance (right) periods while controlling task confounds. (h) Sankey diagram summarizing neurons showing significant MI with latent variables during intertrial (left) and asteroid avoidance (right) periods. (i–k), Tuning curves showing firing rates averaged across trials ranked by latent-variable percentile. Colored shading indicates the low– and high-value ranges used for comparison, defined by the lower 30th and upper 30th percentile ranges (i.e., *<*30th and *>*70th percentiles). Statistics refer to the contrast between these two extremes. (i) Intertrial: uncertainty (Cohen’s *d* = 2.91, *p* = 0.0001), (j) asteroid avoidance: uncertainty (Cohen’s *d* = 2.19, *p* = 0.0001), and (k) asteroid avoidance: prediction error (Cohen’s *d* = 3.89, *p* = 0.0001). (Two-sided permutation test, *p <* 0.001 ***)

All 110 recorded CLA units met the model-fit threshold (*R*^2^ *≥* 0.2) and were included in the latent-variable analyses. Single-neuron analysis revealed that 28.2% of CLA neurons (*n* = 31) were modulated by uncertainty, showing stronger firing in high-uncertainty trials during asteroid avoidance epoch (Fig. 5d top; cluster-based permutation, *p <* 0.05). These neurons also exhibited sustained crash-related firing under high-uncertainty (Fig. 5d, middle) but not low-uncertainty conditions (Fig. 5d, bottom), suggesting that neurons encoding uncertainty-related signals may show enhanced responses to action-guided expectation violations [25] or to salient or aversive outcomes more broadly [3] when the internal beliefs are less precise.

For PE responses, 22.7% of CLA neurons (*n* = 25) showed increased activation under high-PE conditions during the outcome epoch (Fig. 5f top). These neurons exhibited enhanced crash responses with high-PE trials (Fig. 5f middle) but not with low-PE trials (Fig. 5f bottom). Notably, these neurons showed substantial overlap with the uncertainty-modulated population: 23 of 31 uncertainty-modulated CLA neurons were also PE-modulated, and 19 of 25 PE-modulated neurons were also uncertainty-modulated. Because prediction error co-varies with uncertainty over the safety probability, with larger errors typically arising when internal beliefs are less precise (see Extended Data Figure 9a), this overlap is expected and limits the extent to which these signals can be dissociated at the single-neuron level. Although the CMI analysis controlled for shared task confounds, the intrinsic correlation between uncertainty and prediction error means that the present analyses do not establish whether these neurons independently encode uncertainty, PE, or both. We therefore interpret this pattern as indicating that CLA neurons preferentially increase their responses when the internal model is imprecise, particularly when outcomes violate expectations, consistent with a role in prioritizing bottom-up information under conditions of low belief confidence [31].

#### 2.5.1 Conditional Mutual information reveals diverse tuning patterns in CLA

To overcome limitations of quantile-based binarization and potential confounds of co-varying task variables, we used conditional mutual information (CMI) to detect nonlinear relationships between neural activity and model-inferred uncertainty or prediction error [32], while controlling for task-related variables, including spaceship *y*-position, safety-zone identity, and action-contingent outcome (Fig. 5g). For the inter-trial period analyses, we used the period *−*4 to 0 s to ensure that the asteroid from the preceding trial was no longer visible on the screen. Using this time window, 3.6% of neurons (*n* = 4) showed significant MI with uncertainty (Fig. 5h; Extended Data Figure 6c).

During the asteroid-avoidance period (0–1.5 s), the fraction of CLA neurons showing significant MI with uncertainty increased to 9.1% (n=10). Additionally, 6.4% (n=7) of neurons shared significant MI with prediction error (Fig. 5h; Extended Data Figure 6d). These results suggest that CLA neurons contain information about the internal belief-related variables, including uncertainty and prediction error, beyond that explained by external task variables.

Diverse tuning profiles were observed across neurons that shared significant MI with latent variables. Spike counts varied as a function of latent variables during the intertrial and asteroid-avoidance periods. Some neurons increased spike counts at intermediate latent-variable percentiles, while others did so at lower or higher quantiles. Some neurons fired in a monotonic fashion across the ranked variable range, and others showed reduced firing at intermediate values with relatively higher responses at both extremes (Fig. 5i-k; Extended Data Figure 6e-g). These patterns suggest that belief-related latent-variable coding in the CLA is distributed across neurons with heterogeneous tuning profiles for uncertainty and prediction error.

### 2.6 ACC neurons encode uncertainty and PE

We next examined ACC neuronal responses using the same analytical approach. 72 neurons (90% of recorded units) were included in the latent-variable-related analysis; 6 units from Subject 4 and 2 units from Subject 5 were excluded because the corresponding behavioral model fits were low (*R*^2^ *<* 0.2)(Extended Data Figure 3). We identified 23.6% of ACC neurons (*n* = 17) that were responsive to uncertainty. These neurons showed elevated firing under low-uncertainty conditions both during the inter-trial period and during the action-contingent outcome period (Fig. 6b top) but did not distinguish between crash and avoidance outcomes (Fig. 6b middle and bottom), suggesting that they may represent internal model reliability or belief precision [33].

**Fig. 6.**
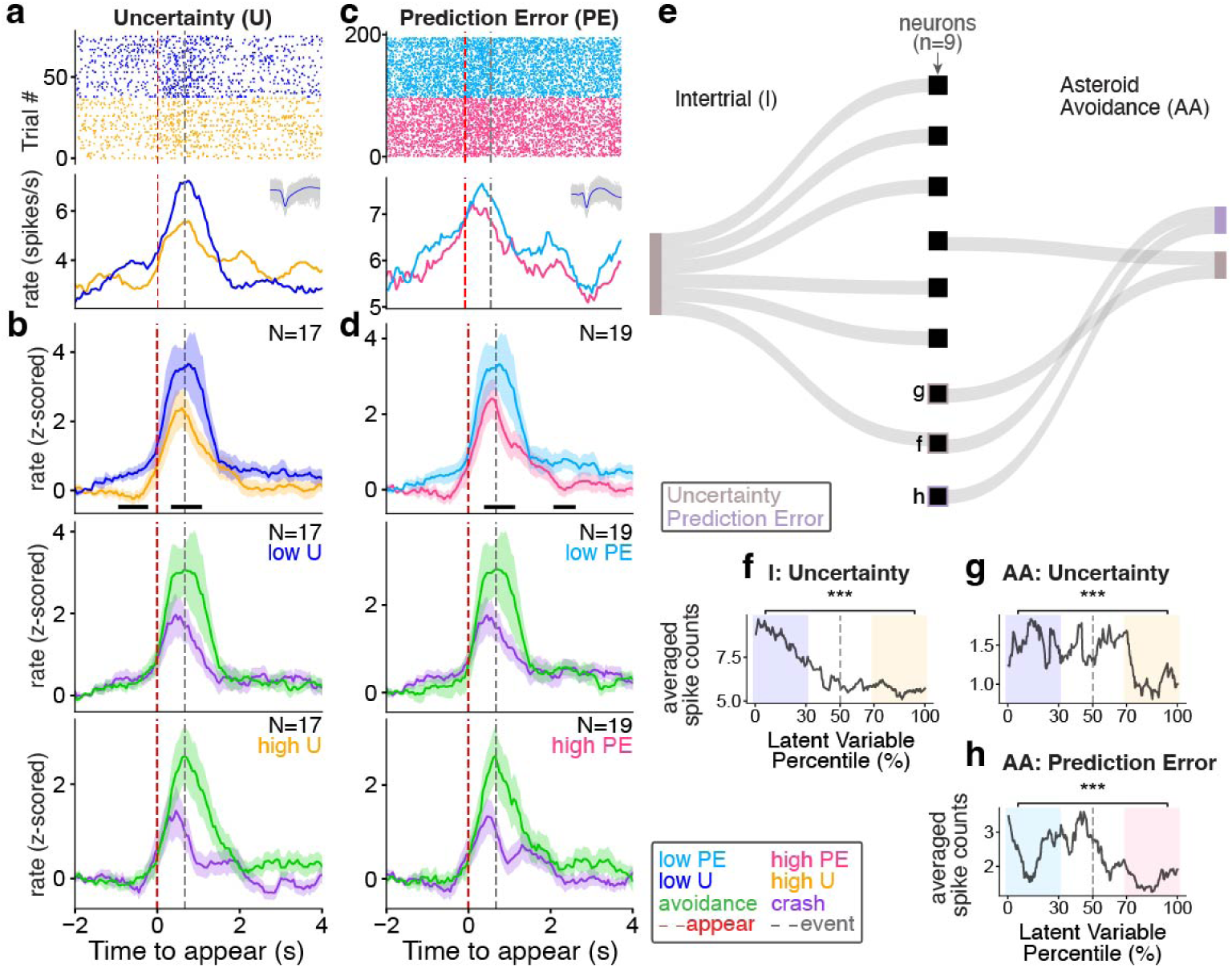
ACC neurons track uncertainty and prediction error. (a,c) Exemplar neurons modulated by uncertainty (a) and (c) prediction error. Neurons were identified based on responses during low-uncertainty or low-prediction-error trials. Top: Spike rasters aligned to task events. Bottom: Mean firing rate (*±* s.e.m.) and waveform of the same neuron. (b, d) Population responses of neurons identified as low-uncertainty–modulated (b; *n* = 17) or low-prediction-error–modulated (d; *n* = 19). Top: Z-scored firing rates (*±* s.e.m.) comparing high versus low uncertainty (b) or high versus low prediction error (d) trials. Middle: Responses during low-uncertainty or low-prediction-error trials, with trials separated by action-contingent outcome (avoidance, green; crash, purple). Bottom: Responses during high-uncertainty or high-prediction-error trials, with trials again separated by outcome (avoidance vs crash). Black bars indicate time periods during which neural responses differed significantly between conditions, identified using two-sided cluster-based permutation test (b) *t* = *−*3.67, *t* = *−*2.98; (d) *t* = *−*3.32, *t* = *−*4.33. (*p <* 0.05). (e) Sankey diagram summarizing neurons showing significant MI with latent variables during intertrial (left; uncertainty) and asteroid avoidance (right; uncertainty and prediction error) periods. (f–h) Tuning curves showing average spike counts across trials ranked by latent-variable percentile. Colored shading indicates the lower and upper extremes of the ranked distribution, defined by the 30th and 70th percentile cutoffs, which were used for statistical comparison. (f) Intertrial: uncertainty (Cohen’s *d* = *−*5.21, *p* = 0.0001), (g) asteroid avoidance: uncertainty (Cohen’s *d* = *−*2.85, *p* = 0.0001), and (h) asteroid avoidance: prediction error (Cohen’s *d* = *−*2.09, *p* = 0.0001). (Two-sided permutation test, *p <* 0.001 ***).

For PE, 19 neurons (26.4%) showed stronger firing during low-PE trials, and did not differentiate between action-contingent outcomes (Fig. 6d). The overlap between these populations was near-complete: all 17 uncertainty-modulated neurons were also PE-modulated, and 17 of 19 PE-modulated neurons were likewise uncertainty-modulated. This convergence further suggests that these ACC neurons may not encode uncertainty and prediction error as fully independent quantities. Rather, they may reflect a composite inferential signal related to belief reliability or the precision-weighted gain with which outcome mismatches drive internal updating, as envisioned in predictive-coding frameworks [22, 23]. These results suggest that responses to belief–outcome mismatches in the ACC are stronger when the internal model is more precise, consistent with precision-weighted processing.

CMI analyses also identified a subset of ACC neurons that encoded latent variables (Fig 6e). Significant MI between a subset of ACC neurons and uncertainty was observed during the intertrial period (8.3% (*n* = 6), Fig. 6f; Extended Data Figure 7c,e). During the asteroid avoidance period, fewer ACC neurons displayed significant MI with uncertainty (2.8% (*n* = 2) or prediction error (2.8% (*n* = 2), Fig. 6g,h; Extended Data Figure 7d,f,g). Compared to CLA, the ACC had a larger fraction of uncertainty-related units during the intertrial period, whereas CLA had a larger fraction of neurons sharing mutual information with uncertainty and prediction error during the asteroid avoidance period.

### 2.7 Amygdala exhibits limited latent variable-related modulation

We next examined amygdala neurons (AMY; Extended Data Figure 10) as a comparison region, given its anatomical proximity to the CLA and its established roles in encoding outcome valence and prediction error [34–37]. A total of 75 AMY neurons were recorded from subjects with simultaneous CLA and ACC recordings.

**Appear-modulated neurons**: Twelve neurons (3 bursters, 9 pausers) displayed significant firing-rate modulation time-locked to asteroid appearance (0.72 *±* 0.30 vs 0.93 *±* 0.38 spikes/s mean *±* s.e.m; *Z* = 2.04, *p* = 0.0425, Wilcoxon signed-rank, *n* = 12; Extended Data Figure 10d right). At the population level, these neurons also differentiated between action-contingent outcomes, with larger absolute firing-rate changes following crash than avoidance outcomes (0.93 *±* 0.35 vs 1.22 *±* 0.44 spikes/s mean *±* s.e.m, *Z* = 2.59, *p* = 0.007, Wilcoxon signed-rank, *n* = 12; Extended Data Figure 10d right).

**Outcome-modulated neurons**: Sixteen neurons (2 bursters, 14 pausers) showed significant firing-rate changes related to action-contingent outcomes (0.47 *±* 0.22 vs 0.78*±*0.28 spikes/s mean *±* s.e.m; *Z* = 3.52, *p* = 0.0001, Wilcoxon signed-rank, *n* = 16; Extended Data Figure 10e right), but did not significantly discriminate between crash and avoidance outcomes at the population level (0.76 *±* 0.27 vs 0.90 *±* 0.30 spikes/s mean *±* s.e.m, *Z* = 2.02, *p* = 0.05, Wilcoxon signed-rank, *n* = 16).

**Appear-and-outcome modulated neurons**: Five neurons (6.7% of units) showed significant modulation in both appearance– and outcome-aligned analyses, indicating responsiveness to both asteroid onset and the subsequent action-contingent outcome; however, this observation is based on a small number of units and should be interpreted cautiously.

Among appear-modulated neurons, firing-rate changes were greater following crash than avoidance outcomes, a pattern consistent with valence-sensitive encoding previously reported in the amygdala [34, 37]. Only 30% of AMY neurons exhibited task-related modulation, compared to CLA (71%) and ACC (70%), and the majority of task-responsive AMY neurons were pausers. Both appearance modulated and outcome modulated neurons were identified; however, significant modulation by outcome type was evident only among appearance-modulated neurons, which show stronger firing with crash compared to avoidance outcomes. Outcome-modulated neurons did not reliably distinguish between crash and avoidance. We also observed minimal latent variable (uncertainty– and prediction error-related) modulation in the amygdala, with only four neurons (one uncertainty-modulated and three prediction error–modulated neurons) meeting inclusion criteria (examples in Extended Data Figure 10f,g). The relatively small proportion of task-modulated AMY units suggests less engagement of this region compared to CLA and ACC during this task, which may in turn account for the correspondingly weak model-derived latent variable modulation observed here.

### 2.8 Latent variable decoding by CLA and ACC neurons

To assess whether uncertainty and prediction error could be decoded from single neuron firing rates, we performed decoding analyses using support vector machine (SVM) classifiers at the single-neuron level and summarized performance across recorded populations. For each neuron, trials were classified as low or high uncertainty or prediction error (30th and 70th percentile cutoffs).

During the intertrial period, neither ACC nor CLA showed significant decoding of uncertainty. By contrast, both uncertainty and prediction error were decodable from activity in both regions during the asteroid-avoidance period. Notably, prediction error could not be decoded during the intertrial period in either region. This is consistent with the task structure as prediction error cannot be computed before asteroid onset. During the asteroid-avoidance period, decoding accuracy was higher than matched shuffle controls for uncertainty and prediction error in both CLA and ACC, but not in AMY (one-sided paired permutation test, *p <* 0.05; Extended Data Figs. Extended Data Figure 6h, Extended Data Figure 7h, Extended Data Figure 10h; Table S4).

Decoding accuracies were modest in absolute terms, as expected for single-neuron decoding analyses that aimed at relating neural activity to cognitive variables that guide behavior [38–40], but they reliably exceeded matched shuffles and generalized across cross-validation splits. Across neurons, confusion matrices were approximately balanced between low/high classes, and decoding gains were driven by units with consistently above-chance performance. Together, these findings indicate that latent-variable information is represented in both the CLA and ACC and can be linearly decoded from single-neuron firing.

## 3 Discussion

In this study, we recorded single neurons from the human claustrum, dorsal anterior cingulate cortex, and amygdala during an aversive learning task that required participants to form and update probabilistic beliefs under uncertainty. We found that CLA and ACC were both heavily engaged, with approximately 70% of neurons in each region showing task-related modulation, compared to only 30% in the AMY. Despite similar overall engagement, the two regions diverged in important ways. CLA neurons exhibited a population-level bias toward crash outcomes, whereas ACC neurons were more heterogeneously tuned, with nearly balanced subpopulations preferring crash and avoidance outcomes. Both regions encoded model-derived latent variables — uncertainty and prediction error — but with different temporal and functional profiles: ACC neurons carried uncertainty-related signals during the intertrial period, consistent with a role in maintaining belief precision prior to new evidence, while CLA neurons predominantly showed latent-variable encoding during the asteroid-avoidance period, when prediction errors could be computed and salient outcomes demand rapid updating. Conditional mutual information analyses confirmed that these signals were present beyond what could be explained by external task variables. Single-neuron decoding demonstrated that latent-variable information was linearly accessible from firing rates in both regions, but not in the AMY. Together, these findings suggest complementary roles for the CLA and ACC in tracking latent task states within a predictive processing framework: the ACC appears to maintain and monitor internal model reliability, while the CLA preferentially signals when outcomes violate goal-directed expectations under conditions of low model confidence — a pattern consistent with the release of bottom-up error signals from top-down suppression when belief precision is low, potentially facilitating coordinated cortical updating in dynamic environments.

Our recordings demonstrate that CLA neurons are sensitive to latent task variables, suggesting a role in hierarchical inference [24, 41]. Macaque anatomical-tracing studies [5] support this view: the CLA contains projection-specific zones that integrate convergent inputs from prefrontal, cingulate, sensory, and entorhinal cortices while producing topographically divergent outputs — an architecture well suited to context-sensitive transformations between distributed cortical computations. Rodent studies further demonstrate that the CLA determines whether prediction–input mismatches are dismissed as noise or propagated as meaningful signals, indicating context-dependent sensory gain regulation [42]. Notably, disorders involving sensory-perceptual disturbances — including autism, schizophrenia, and psychedelic states — consistently show CLA structural or functional abnormalities[3]. Mechanistic work in mice has shown that inhibiting CLA*→* prefrontal projections increases resting frontal and sensory-evoked visual cortical activity, while activation suppresses prefrontal output [43], supporting the CLA’s proposed inhibitory role in cortical dynamics [8]. ACC inputs convey load-proportional preparatory signals to the CLA, where local microcircuits amplify and constrain them in a frequency-dependent manner [12]. Our findings extend this framework to humans, suggesting that the CLA, by encoding latent task structure, contributes to context-sensitive neural resource allocation and may support flexible top-down control adjustment based on inferred task demands, as evidenced by enhanced responses to crash outcomes under heightened uncertainty or prediction error.

In contrast, ACC neurons predominantly showed increased activation during low-uncertainty or low-prediction-error conditions. This pattern differs from prior primate single-unit reports in which ACC firing rates scaled positively with uncertainty [18] and prediction error [44], as well as from fMRI studies showing that BOLD signal in ACC increases with greater uncertainty [28], prediction error [45], or belief updating [46]. These discrepancies may arise from several factors. First, latent-variable modulation is task-dependent: the prediction error in this study is defined as a belief–observation mismatch within a Bayesian framework, which differs from prior ACC studies that have operationalized prediction error in the context of action–outcome expectation violations[19, 29, 47–51]. Within predictive coding theory, neural responses are often interpreted as reflecting precision-weighted prediction errors, such that stronger activity under low uncertainty may indicate increased gain on prediction errors when internal beliefs are precise [21–23, 52, 53]. Second, BOLD signals reflect aggregated population activity and are not a direct measure of single-neuron firing rates; quantitative fMRI studies have shown that ΔBOLD may, in certain contexts, oppose metabolic measures, indicating that BOLD is not a sign-preserving index of neuronal activity [54]. Third, anatomical sampling may also be relevant, as the ACC exhibits marked subregional heterogeneity [55]. We primarily recorded from areas 24a/24b, whereas prior studies have often examined dorsal sulcal regions (24c/32) [18, 29, 48]. Taken together, these data suggest that a subset of ACC neurons encode model-derived latent variables in a manner consistent with belief-precision signaling.

Although our study was not designed to test causal interactions between regions, the complementary temporal profiles we observed — ACC encoding of uncertainty during the intertrial period and CLA encoding of latent variables during the asteroid-avoidance period — raise the possibility that these regions may operate as part of a coordinated circuit rather than independently. Anatomical evidence supports this interpretation: the ACC sends direct projections to the CLA, and prior work in rodents has shown that these inputs convey load-proportional preparatory signals that are locally amplified within claustral microcircuits [12]. One speculative model is that ACC precision signals maintained during the intertrial interval normally exert a tonic suppressive influence on CLA output, consistent with the inhibitory role of ACC*→*CLA projections described in rodents [8, 43]. When uncertainty is high, this precision-related input weakens, effectively releasing the CLA from top-down constraint and allowing stronger propagation of bottom-up prediction-error signals during the subsequent avoidance period. This interpretation aligns with our observation that CLA neurons showed enhanced crash responses specifically under high-uncertainty conditions: reduced ACC-mediated suppression would permit greater CLA responsiveness to salient outcome mismatches precisely when the internal model is least certain. More broadly, this account is consistent with predictive processing frameworks in which precision estimates from higher-order regions gate the propagation of prediction errors in downstream circuits — not by amplifying errors under certainty, but by suppressing them, such that errors are propagated more freely when precision is low [21, 23]. However, establishing whether such a functional relay exists will require simultaneous multi-region recordings with sufficient spatial and temporal resolution to assess directed information flow, or causal perturbation approaches such as targeted microstimulation of ACC*→*CLA projections.

A large body of work has reported prediction-error or surprise signals in the amygdala [35, 37, 56]. However, our recorded AMY neurons showed little evidence of latent-variable encoding. This discrepancy may reflect, at least in part, differences in the computational variables examined. Whereas most prior studies quantify amygdala responses relative to outcome–expectancy violations, our analyses tested for encoding of latent belief-state variables derived from a Bayesian framework — a distinction for which direct single-unit evidence in the amygdala is lacking [57]. Convergent evidence suggests that the amygdala’s learning-related signals primarily reflect quantities that modulate the strength of learning, such as associability or surprise, rather than variables that encode inferred latent states of the environment [57–60]. This dissociation has been demonstrated in humans, where bilateral amygdala BOLD activity tracks associability signals that are dissociable from ventral striatal responses to outcome–expectation discrepancies during aversive learning [61]. Nevertheless, the absence of Bayesian belief-related modulation in our data should be interpreted cautiously given the limited number of units recorded from human subjects.

Several limitations warrant consideration. CLA responses may partially reflect sensory confounds. Our epilepsy patient cohort, potential effects of antiseizure medications, and the pseudo-population analytical approach may limit generalizability; our conclusions pertain to the sampled neuronal populations rather than to complete regional repertoires. Latent-state inference relied on an approximate Bayesian updating model validated in large datasets [2], chosen for its capacity to capture trial-by-trial evidence accumulation and posterior uncertainty, not to imply that the brain implements this specific algorithm. The model tests whether neurons represent abstract task variables without committing to a particular computational formalism.

Future work could employ microstimulation of CLA neural populations to causally dissect their role in inferential dynamics. A key question is how transient inhibition of the CLA influences behavior and the trial-wise dynamics of latent cognitive variables. In parallel, it will be important to assess whether the CLA contributes to the modulation of precision-weighted prediction errors across the cortex. This could be tested by measuring changes in the trial-wise magnitude and variability of prediction-error signals in sensory and prefrontal areas before and after claustral suppression. Answering these questions would clarify whether the CLA functions as a modulatory hub for hierarchical inference and belief-guided sensory gating.

## 4 Methods

### 4.1 Participants and sessions

Seven subjects (2 males, 5 females; ages 18-61; Table S1) with medication-refractory epilepsy underwent intracranial monitoring for seizure onset localization at the Yale Comprehensive Epilepsy Center. Hybrid depth electrodes with 8 microwires were implanted to record single-unit activity during task performance. Electrode placement was determined clinically. Subjects with seizure onset or interictal epileptic discharges in claustrum, anterior midcingulate cortex or amygdala electrodes were excluded. All subjects provided informed consent under research protocols approved by the Yale School of Medicine Institutional Review Board.

### 4.2 Behavioral task

Subjects performed a validated aversive learning task [2], on a computer (800 *×* 1600 pixels; Fig. 1c), controlling a spaceship along the y-axis to avoid asteroid collisions and maximize points. Spaceship integrity and score were displayed onscreen. Crashes decreased spaceship integrity while successful avoidance restored it; if integrity reached zero, the game ended, though participants could resume. Safety holes appeared in either the top or bottom asteroid belt, with safety probabilities varying independently and dynamically across trials. Safety outcomes were statistically independent, preventing cross-zone inference. Successful performance required anticipating the likely safe zone and positioning the spaceship before asteroid appearance, as post-asteroid appearance avoidance was rarely achievable.

The task comprised blocks of 20 and 80 trials, with one safety zone assigned 90% or 10% safety probability, reversing randomly across blocks. Each trial guaranteed at least one safe zone, ensuring avoidance remained possible while maintaining environmental instability. A vigilance check required button presses to sustain attentional engagement.

Behavioral data were analyzed offline with variable sampling rates (50–90 samples/second). Task variables included spaceship *y*-position, asteroid appearance, spaceship integrity, and behavioral outcomes. An approximate Bayesian model inferred subjects’ internal states, including uncertainty and prediction error per trial (see “Asymmetric Leaky Beta Model” below). For asteroid onset and outcome responses, we extracted asteroid appearance times and identified moments of sharp spaceship integrity decrease. Post-event movement was measured over 3-second windows. For successful avoidance responses, we estimated average time from asteroid appearance to first crash (mean: 645ms, Fig. 1e) to define avoidance timing. Each subject completed 1-2 sessions; data from separate days (Subjects 1 and 3) were treated as independent datasets (Table S1).

#### Asymmetric leaky beta (ALB) model

The ALB model is an approximately Bayesian model that uses beta distributions to represent an approximate distribution over safety probabilities (note that this is approximate since the full posterior distribution is not derived exactly). The model, previously employed to characterize aversive learning in controlled laboratory tasks [62] and unsupervised online experiment platforms [2], conceptualizes agents as estimating beta distributions representing the probability of encountering a safe event (e.g., presence of a hole in the asteroid belt) independently for each location. The beta distribution, a continuous probability distribution bounded between 0 and 1, is particularly suited for representing uncertainty about probabilities [63], with its shape determined by two positive parameters: count of positive outcomes (e.g. avoid the asteroid) and count of negative outcomes (e.g. crash the spaceship). The “leak” in the model is crucial for learning non-stationary outcome probabilities, as it allows estimates of safety probability to decay over time, ensuring that the agent’s behavior is most strongly influenced by recent outcomes. The asymmetric component enables the model to capture well-established valence asymmetries in learning, where participants tend to exhibit enhanced learning for danger over safety in aversive learning tasks [2, 62].

#### Model Characteristics and Parameters

- **Asymmetric**: separate learning rates for positive (safe, *τ* ^+^) and negative (dangerous, *τ ^−^*) outcomes.
- **Leaky** (*λ*): allows accumulated evidence to decay over time, allowing adaptation in volatile environments.
- **Probabilistic Representation**: captures both mean and variance of the estimated safety.

#### Model Updates

For each location (A or B), trial outcomes update two accumulators:

*α*: evidence for safety.
*β*: evidence for danger.

Discrete trial outcomes (safe = 1, danger = 0) update these accumulators additively by respective update rates for safe and dangerous outcomes. Specifically, updates at trial are given by:

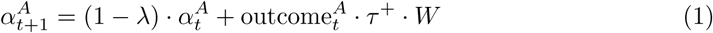

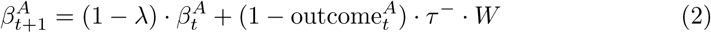

where attention weight parameter is defined as:

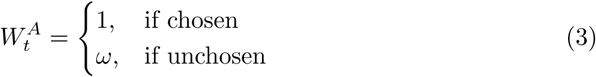

and enables the model to capture preferential attention to chosen location.

#### Estimating Safety Probability and Uncertainty

The model calculates the probability of safety at each location (e.g. A-Top) from the mean of the current beta distribution on each trial:

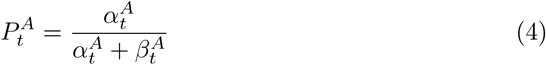

The associated uncertainty *σ*, represented as the variance of the beta distribution, is computed by:

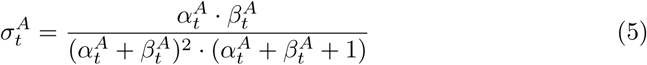

#### Behavioral Response and Choice Stickiness

Subjects moved toward safer locations, adopting neutral positions under uncertainty (Fig 5a). A stickiness parameter accounts for habitual choice repetition, increasing the likelihood of selecting the previously chosen location regardless of outcome:

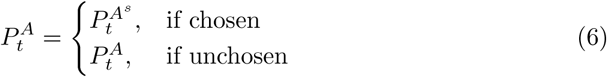

#### Estimating Prediction Error

Prediction error (*δ*) is defined as the difference between the actual trial outcome and the model-predicted safety probability for a given location. We focused on the absolute magnitude of the prediction error to quantify surprise, regardless of outcome valence, as a measure of expectation violation modulating neural activity. Absolute prediction error is defined as:

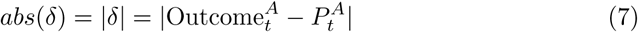

#### Determining Behavioral Position

The spaceship’s position is guided by predicted safety probabilities, shifting toward the safer location. If both options are equally uncertain, the position remains near the center.

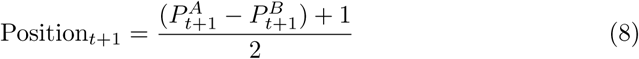

#### Model Fitting

The same model-fitting approach as in the previous study was applied, using variational inference to estimate posterior distributions over model parameters [2]. The mean of this distribution was then taken as a point estimate for each parameter for each subject, and simulated data (providing mean and uncertainty estimates, along with prediction errors) were generated using the model with parameters set to these estimated values. To quantify goodness-of-fit, we computed subject-wise *R*^2^ values, reflecting the proportion of variance in observed behavior accounted for by the fitted model. Higher *R*^2^ values therefore indicate closer agreement between model-predicted and empirical behavior at the individual-subject level (Extended Data Figure 3).

For completeness, and to benchmark our approach against models commonly used in prior studies, we also tested two reinforcement learning accounts of behavior: a standard Rescorla–Wagner model and a variant with dual learning rates [64]. Consistent with a previous large-sample study [2], our model comparison showed that the ALB-sticky model provided the best fit to the behavioral data (see Extended Data Figure 3).

### 4.3 Robotic-assisted surgical implantation of intracranial depth electrodes

Behnke-Fried depth electrodes (AD-TECH®, Wisconsin, USA; 8 microwires per macroelectrode) were implanted to map the seizure network, as determined by the clinical team. Electrode placement was planned using preoperative T1-contrast MRI fused with high-resolution CT scans via the ROSA One®robotic platform (Zimmer Biomet; 0.3mm accuracy). Claustral microwires were positioned in depth electrodes targeting the middle insula, with macroelectrode contacts in the insular cortex and only microwires extending into the claustrum (see Extended Data Figure 1). No neurovascular injuries were observed.

### 4.4 Electrode localization

Pre-operative T1-weighted MRI sequences and post-operative CT scans were co-registered using LeGUI [65] (Extended Data Figure 1, Extended Data Figure 2). Two expert neurosurgeons visually verified microwire locations. Electrode and microwire positions were normalized to MNI space within LeGUI (Table S2) and visualized in 2D/3D using MRIcroGL software (https://www.nitrc.org/projects/mricrogl). The position of microwires targeting the claustrum was also confirmed in the MNI space using Lead-DBS [66], which uses Advanced Normalization Tools (ANTs) [67] for CT–MRI co-registration and nonlinear normalization prior to manual inspection of electrode locations.

### 4.5 Electrophysiological recordings

Electrophysiological data were recorded during the task using Neuroport system (Blackrock Microsystems®, Salt Lake City, Utah USA) at 30kHz with 0.3Hz-7.5kHz hardware filter. One microwire per electrode served as local reference. Data were stored and analyzed offline.

### 4.6 Spike detection and sorting

Raw signals were filtered with a zero-lag phase filter from 300-3000Hz and spike-sorted using an automated template-matching algorithm [68], considering both positive and negative deflections. Single units were defined by template matching and analyzed units were included if the following criteria were met: at least 30 trials per condition (crash or avoidance), the isolated waveform visually matched the profile of an action potential, inter-spike interval (ISI) of *<* 3ms was present in *<* 5% of the detected spikes, Poisson-distributed ISI, signal to noise ratio (SNR) *>* 1, and stable recording throughout the recording of the task. Multiunits were excluded from the analyses.

### 4.7 Stability assessment of single units

Single-unit stability was assessed following prior work [69], by evaluating both temporal consistency of firing and stability of spike waveforms. Temporal firing stability: Cumulative spike distribution was compared to an ideal uniform distribution using root mean squared error (RMSE). Firing rate vs. ISI violations (%) were also evaluated. Waveform stability: From 1000 randomly sampled spikes per unit, we computed five metrics: L-ratio (cluster quality), Coefficient of variation (CoV) of maximum amplitude, CoV of full-width at half-maximum, CoV of waveform Area Under the Curve, Euclidean distance from the mean waveform. Each of these metrics were computed for each unit, and their CoV was calculated across the repeated random subsampling (Extended Data Figure 8).

### 4.8 Single unit analysis

Peri-stimulus time histograms (PSTHs) were computed using 50ms bins aligned to asteroid appearance or outcomes (crash/avoidance). Trial-averaged firing rates were used for population analysis (see below). Raster plots and 200ms sliding window mean firing rates across trials were calculated for example neurons.

### 4.9 Appear and outcome-modulated neurons

To identify task-modulated neurons, we used a two-sided permutation test (*n* = 10, 000 permutations; accessed via MathWorks File Exchange (https://github.com/lrkrol/permutationTest; retrieved 27 September 2025) comparing mean firing rates during an ‘appear window’ (0-500ms after asteroid onset, aligned to asteroid appearance; appear-modulated neurons) or ‘outcome window’ (0-500ms following action-contingent outcome (crash/avoidance), aligned to outcome onset; outcome-modulated neurons) to a baseline (−2000 to –1500ms before asteroid onset, aligned to asteroid appearance). To correct for multiple comparisons across neurons, Benjamini-Hochberg False Discovery Rate (FDR) correction [70] was applied separately to the *p*-values from the appear and outcome windows conditions to control the expected proportion of false positives among all statistically significant (*p <* 0.05) neurons.

Neurons with significant modulation (*p <* 0.05) were classified as appear-, outcome-, or task-modulated. ‘Task-modulated neurons’ were defined as having significant responses to appear or any outcome. Appear-modulated neurons were responsive to asteroid onset regardless of outcome responsiveness. Appear-specific neurons only showed significance during ‘appear window’, while appear-and-outcome neurons showed significance during the ‘appear window’ and ‘outcome window’. Outcome-specific neurons were defined as having a significant response during ‘outcome window’ (i.e., following avoidance or crash), but not to the appear window. Custom MATLAB codes were used to generate Venn diagrams (MATLAB Central File Exchange, Darik 2023, v.1.7, https://www.mathworks.com/matlabcentral/fileexchange/22282-venn).

For analysis comparing CLA and ACC outcome firing patterns (Fig. 4k-o), neurons were categorized based on having a significant response to crash (‘crash-responsive’) or to avoidance (‘avoidance-responsive’), regardless of their response to appearance or to the alternative outcome. This approach avoided circularity in subsequent population comparisons between crash and avoidance responses.

Percentages of responsive neurons (Fig. 2) were calculated using a permutation test comparing mean firing rates to the neuron’s baseline per time bin. To correct for multiple comparisons across neurons, Benjamini-Hochberg False Discovery Rate (FDR) correction was applied on every *p*-value obtained per time bin. Neurons with *p <* 0.05 after FDR correction were considered responsive.

### 4.10 Average population analysis

Task-modulated neurons were pooled by brain region. Firing rates were z-scored relative to the baseline window and averaged (50ms bins; –2 to 4 s relative to event onset). When the baseline mean firing rate for a neuron was zero, normalization was per-formed using the mean firing rate across the full trial rather than the baseline alone. For line plots showing mean population responses (Fig. 3e,f,h,j, Fig. 4e,g,i,k-m, and Extended Data Figure 10d,e), z-scored firing rates were smoothed (200ms) and averaged. For heatmaps (Fig. 3d and Fig. Extended Data Figure 10a), absolute changes from baseline were normalized (0= no change, 1= maximum change) and sorted by latency of maximum response after asteroid onset (early on the bottom). For heatmaps showing multiple neurons (Fig. 3e, Fig. 4d, and Extended Data Figure 10b), firing rates of appear-modulated units were normalized (0 to 1) and sorted based on their mean rate (low at the bottom) during the ‘appear window’. Change in firing rate (Δ firing rates) was assessed by subtracting baseline firing rate from mean firing during the ‘outcome window’ (crash or avoidance; Fig. 3g,i,k; Fig. 4f,h,n,o; Extended Data Figure 10d,e). For appear– and outcome-modulated neurons (Fig. 3h,i; Fig. 4g,h,k,l,m; Extended Data Figure 10d,e), absolute firing rate changes were computed by subtracting the mean rate in the ‘appear’ or ‘outcome window’ from baseline. Four windows were defined: pre-appear (−500 to –50ms), post-appear (0 to 500ms), post-avoid (0 to 500ms), and post-crash (0 to 500ms).

### 4.11 Linear mixed-effects model for outcome and region effects

Firing-rate data from task-modulated neurons were analyzed using a linear mixed-effects model with action-contingent outcome (avoidance, crash), region (CLA, ACC), and their interaction as fixed effects. Random intercepts were included for neuron, to account for repeated measurements across outcomes within the same neuron, and for subject, to account for non-independence among neurons recorded from the same participant. Models were fit by maximum likelihood using pandas 2.2.3 and statsmodels 0.14.6.

To evaluate the robustness of the linear mixed-effects results, we performed two additional sensitivity analyses. First, we carried out a leave-one-subject-out procedure in which the same model was refit repeatedly after excluding each subject in turn, allowing us to determine whether the region *×* outcome interaction was disproportionately influenced by any individual participant. Second, to address the non-normal and right-skewed distribution of firing rates, we repeated the analysis after applying a log transformation to firing rates, log(1 + fr), and refit the identical model structure to test whether the interaction remained stable under a reduced-skew representation of the data.

### 4.12 Sign test for neuron-wise outcome preference

To quantify heterogeneity in outcome tuning across neurons, we computed a neuron-wise outcome preference index for each task-responsive unit as the difference between its mean firing rate on avoidance and crash trials (Δ = FR_avoidance_ *−*FR_crash_). Neurons were then classified according to the sign of this difference as avoidance-preferring (Δ *>* 0), crash-preferring (Δ *<* 0), or non-selective (Δ = 0). To test whether one preference direction was overrepresented within a region, we applied a two-sided exact binomial sign test to the counts of positive versus negative Δ values, excluding neurons with Δ = 0. This analysis was used to assess whether the population showed a consistent directional bias or instead exhibited heterogeneous mixed-sign tuning across units.

### 4.13 Neurons modulated by prediction error and subjective uncertainty

Prediction error, subjective uncertainty, and safety probability were estimated using the ALB-sticky model, which provided trial-wise scores for each latent variable in zone A and zone B. For prediction error (PE) and uncertainty (U), we averaged zone A and zone B values within each trial to obtain a single trial-wise value (i.e., 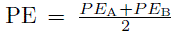 and 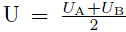), which captures an integrated estimate of prediction error/uncertainty across the two simultaneously tracked locations and avoids duplicating population analyses. Trials were binarized into “low” (*≤* 30th percentile) and “high” (*≥* 70th percentile) (Python quantile functions), requiring *≥* 30 trials per group; neurons with fewer trials were excluded.

For each neuron and variable split (low and high trials), firing rates were demeaned per trial by subtracting the mean firing of a baseline window (−2.0 to –1.5 s before asteroid onset), yielding baseline-corrected responses. We then identified neurons with significant post-stimulus increases in firing relative to baseline using a one-sided cluster-based permutation test. This selection was performed separately for high and low groups, focusing on response presence rather than differential tuning, thereby avoiding circularity in subsequent between-group comparisons (*p <* 0.05, *n* = 10,000 shuffles). Time periods during which firing rates differed significantly between conditions (identified using a two-sided cluster-based permutation test (*p <* 0.05)) were deemed significant, and a neuron was labeled “High-modulated” or “Low-modulated” if the corresponding high– or low-condition test yielded at least one contiguous time window of significant difference lasting *≥* 100ms.

To assess latent-variable effects at the population level, we computed each neuron’s peri-event time course by averaging firing rates across trials within each condition (e.g., for “High-modulated” neurons, each row contained the neuron’s mean high-trial and companion low-trial means). These neuron-level mean time courses were then normalized by z-scoring relative to the same baseline window (−2.0 to –1.5 s before asteroid onset). This z-scoring step was applied only to the neuron-averaged time courses used for visualization and was not applied in addition to the trial-wise baseline subtraction used for the permutation tests. For visualization, firing rates were smoothed with a 400 ms sliding window, a kernel size commonly used to stabilize time courses for cluster-based permutation testing while retaining cognitive-scale dynamics and improving interpretability [71]. Population-level comparisons of High vs. Low and Crash vs. Avoidance conditions were conducted using a paired two-sided cluster-based permutation test across neurons (*n* = 10,000 shuffles). This approach enabled the detection of significant differences in either direction (*p <* 0.05), identifying populations showing both increases and decreases in firing rates between conditions. Observed *t*-statistics were computed at each time bin and thresholded using the critical *t*-value for *p <* 0.05 to define contiguous temporal clusters. Cluster mass was defined as the sum of t-values within each cluster. Observed cluster masses were then compared against a permutation-derived null distribution formed from the largest cluster mass obtained in each shuffle, and significant clusters (*p <* 0.05) were plotted as horizontal bars beneath the mean firing rate traces (Fig. 5d,f, 6b,d, Extended Data Figure 6a-b, Extended Data Figure 7a-b).

### 4.14 Conditional Mutual Information Analysis Controlling for Task Confounds

To control for potential task confounds, we quantified the dependence between trial-wise spike-count responses *R* and each latent variable of interest *X* while conditioning on confounds *Y* using a conditional mutual information (CMI) framework (Fig. 5g). Firing rates were aligned to asteroid onset and analyzed from *−*4 to 2 s, using an intertrial analysis window (*−*4 to 0 s) and an asteroid-avoidance epoch (0 to 1.5 s), with 1.5 to 2 s as baseline. For each neuron and trial, spike counts were obtained by summing spikes within each epoch. Prediction error and uncertainty were reduced to a single scalar per trial by taking the mean of the model-derived values for zones A and B.

CMI quantifies the dependence between *R* and *X* that cannot be explained by *Y*:

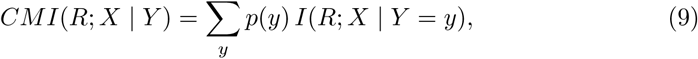

where *I*(*R*; *X | Y* = *y*) denotes the mutual information between *R* and *X* computed across trials with the same confound condition *y*, and *p*(*y*) is the proportion of trials in that condition. Because we used a discrete mutual information estimator (mutual info score, scikit-learn, Python), latent variables *X* were discretized into 10 quantile bins to form discrete labels suitable for discrete mutual information estimation. Confounding variables were discretized as required: spaceship *y*-position was binned into 20-pixel intervals, and categorical variables (Action-contingent Outcome and Safety Belt presence) were treated as discrete labels. In practice, conditional mutual information was estimated by computing mutual information between *R* and *X* separately within trial subsets (strata) sharing the same confound values and averaging these values across subsets, weighted by the number of trials in each subset.

Strata with fewer than three trials (*n_s_ <* 3) were excluded to ensure stable estimation. Confounds differed by epoch, reflecting the task structure. For the intertrial epoch we conditioned on spaceship *y*-position. For the asteroid-avoidance epoch we conditioned on spaceship *y*-position, Safety-Belt presence, and Action-contingent Outcome.

Statistical significance was assessed using a conditional permutation test that preserves the confound structure: within each confound-defined stratum, trial labels for *X* were randomly permuted while holding *R* and *Y* fixed, generating a null distribution of conditional mutual information under the null hypothesis of conditional independence, and neurons with *p <* 0.05 (*n* = 10,000 shuffles) were considered significant (Extended Data Figure 6c,d for CLA; Extended Data Figure 7c,d for ACC). Because prediction error and uncertainty are not statistically independent and our goal was to classify neurons according to the latent variable that best explained their activity rather than to perform independent hypothesis tests for each variable, no additional multiple-comparison corrections were applied [72].

To examine neuronal tuning to latent variables during the intertrial and asteroid-avoidance periods, we compared low versus high latent-variable conditions (trials with percentile rank *≤* 0.30 and *≥* 0.70, respectively). Trial ranks were converted to per-centiles (scipy.stats.rankdata, SciPy, Python), and smoothed firing-rate tuning curves were generated using a symmetric 40-trial moving window (Fig. 5i-k; Fig. 6f-h; Extended Data Figure 6e-g; Extended Data Figure 7e-g; Tables S3). For each neuron, we computed the mean smoothed firing rate across tuning-curve bins corresponding to the low and high percentile ranges of the variable and assessed significance using a two-sided permutation test (*n* = 10,000 permutations). Effect size was quantified with Cohen’s *d*.

### 4.15 Decoding analysis using a linear support vector machine

To assess whether neural activity contained linearly accessible information about task-relevant latent variables, we performed a decoding analysis using a linear support vector machine (SVM). This analysis complements the conditional mutual information approach by testing whether neural activity could discriminate latent variables.

Averaged prediction error and uncertainty across the two spatial zones were used. For each neuron, trials were labeled according to the trial-wise value of the latent variable (uncertainty or prediction error) and binarized into low and high conditions using the 30th and 70th percentiles; intermediate trials were excluded. Firing rates were extracted separately for the intertrial analysis window (*−*4 to 0 s) and the asteroid-avoidance epoch (0 to 1.5 s).

Within each epoch, neural activity was represented as binned firing-rate vectors in 50-ms bins, which were averaged into consecutive non-overlapping 500-ms windows (ten 50-ms bins per window) to reduce feature dimensionality and improve robustness to trial-to-trial spiking variability [73, 74].

Classification was performed using a linear SVM with balanced class weights [75–77] and z-score–normalized features (sklearn.svm.SVC, scikit-learn, Python). A linear kernel was chosen to restrict decoding to information linearly accessible from neural activity. Performance was evaluated with repeated stratified shuffle–split cross-validation (1000 repetitions; 80% training, 20% testing). Accuracy was computed on the held-out set for each split and averaged across repetitions for each neuron.

Statistical significance was assessed using a matched shuffle control in which class labels were randomly permuted within the training set for each split prior to model fitting. True and shuffled accuracies were compared using a one-sided paired permutation test using random sign flips (*n* = 10,000 permutations) across cross-validation splits, testing whether decoding performance exceeded the shuffled control. Effect sizes were quantified using paired Cohen’s *d_z_*. To control for multiple comparisons across neurons, *p*-values were FDR-corrected (Benjamini–Hochberg; *α* = 0.05) separately for each epoch and latent variable. Results are reported as mean decoding accuracy with bootstrap-derived 95% confidence intervals across neurons, computed separately for each brain region, epoch, and latent variable (Extended Data Figure 6h, Extended Data Figure 7h, Extended Data Figure 10h; Table S4).

### 4.16 Neuronal responses following spaceship displacement

To assess if neuronal responses to asteroid appearance, outcomes, or latent variables were driven by predictable visual sensory inputs, activity of appearance-responsive neurons was aligned to spaceship movement onset during the intertrial period (no asteroids present; Extended Data Figure 4). Trials were classified as movement or stationary based on spaceship movement within the 4 s intertrial period, and mean firing rates were calculated per region.

### 4.17 Statistics

Statistical analyses were performed in MATLAB R2023b (MathWorks) and Python 3.9 (packages: NumPy, scikit-learn, statsmodels, SciPy). Population averages were computed as the arithmetic mean, with error bars representing the standard error of the mean (s.e.m.). Non-parametric tests (Wilcoxon signed-rank, rank-sum, Kruskal–Wallis, permutation test, and cluster-based permutation test) were used when data were not normally distributed or involved paired comparisons. Correction for multiple comparisons was performed using the Benjamini–Hochberg false discovery rate (FDR) procedure where noted. Significance thresholds are indicated in the figures (*p <* 0.05, corrected when applicable). Z-scoring was applied for visualization purposes only; all statistical comparisons were performed on Δ firing rates. A complete summary of all statistical tests is provided in Table S5.

## Code and Data availability

Analysis code to reproduce main figures is available at https://github.com/damisahlab/claustrum-uncertainty-surprise. The data generated in this study may be obtained by request to the corresponding author via email.

## Acknowledgements

We thank the members of the Yale Comprehensive Epilepsy Center for their excellent patient care, the patients who participated in this study. This study was supported by grants from the National Institutes of Health (R01MH138291 ECD); The Hypothesis Fund Seed Grant (ECD), VA National Center for PTSD and Wellcome Trust Grant: 310142/Z/24/Z (ECD, AK, and JK).

## Supplementary Information

**Table 1.**
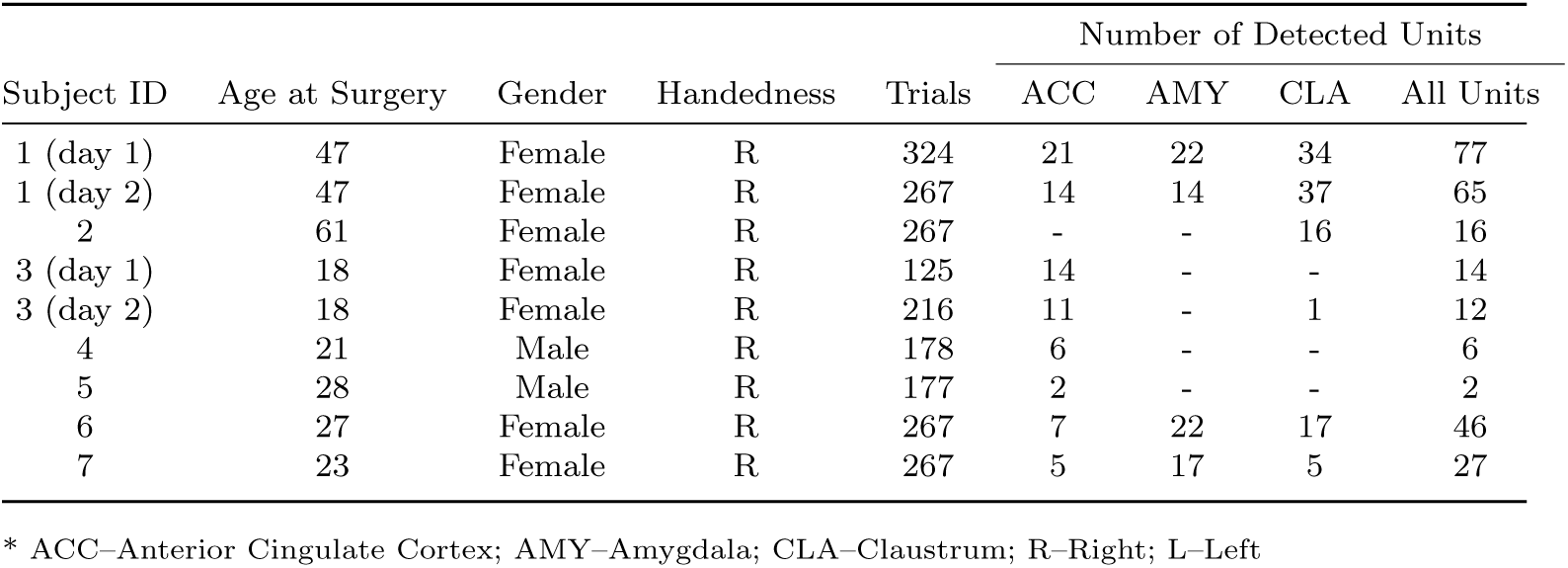
Subject information, trial number, and detected units per region.

| Subject ID | Age at Surgery | Gender | Handedness | Trials | Number of Detected Units |  |  |  |
| --- | --- | --- | --- | --- | --- | --- | --- | --- |
|  |  |  |  |  | ACC | AMY | CLA | All Units |
| 1 (day 1) | 47 | Female | R | 324 | 21 | 22 | 34 | 77 |
| 1 (day 2) | 47 | Female | R | 267 | 14 | 14 | 37 | 65 |
| 2 | 61 | Female | R | 267 | - | - | 16 | 16 |
| 3 (day 1) | 18 | Female | R | 125 | 14 | - | - | 14 |
| 3 (day 2) | 18 | Female | R | 216 | 11 | - | 1 | 12 |
| 4 | 21 | Male | R | 178 | 6 | - | - | 6 |
| 5 | 28 | Male | R | 177 | 2 | - | - | 2 |
| 6 | 27 | Female | R | 267 | 7 | 22 | 17 | 46 |
| 7 | 23 | Female | R | 267 | 5 | 17 | 5 | 27 |

\* ACC–Anterior Cingulate Cortex; AMY–Amygdala; CLA–Caudate; R–Right; L–Left

**Table 2.**
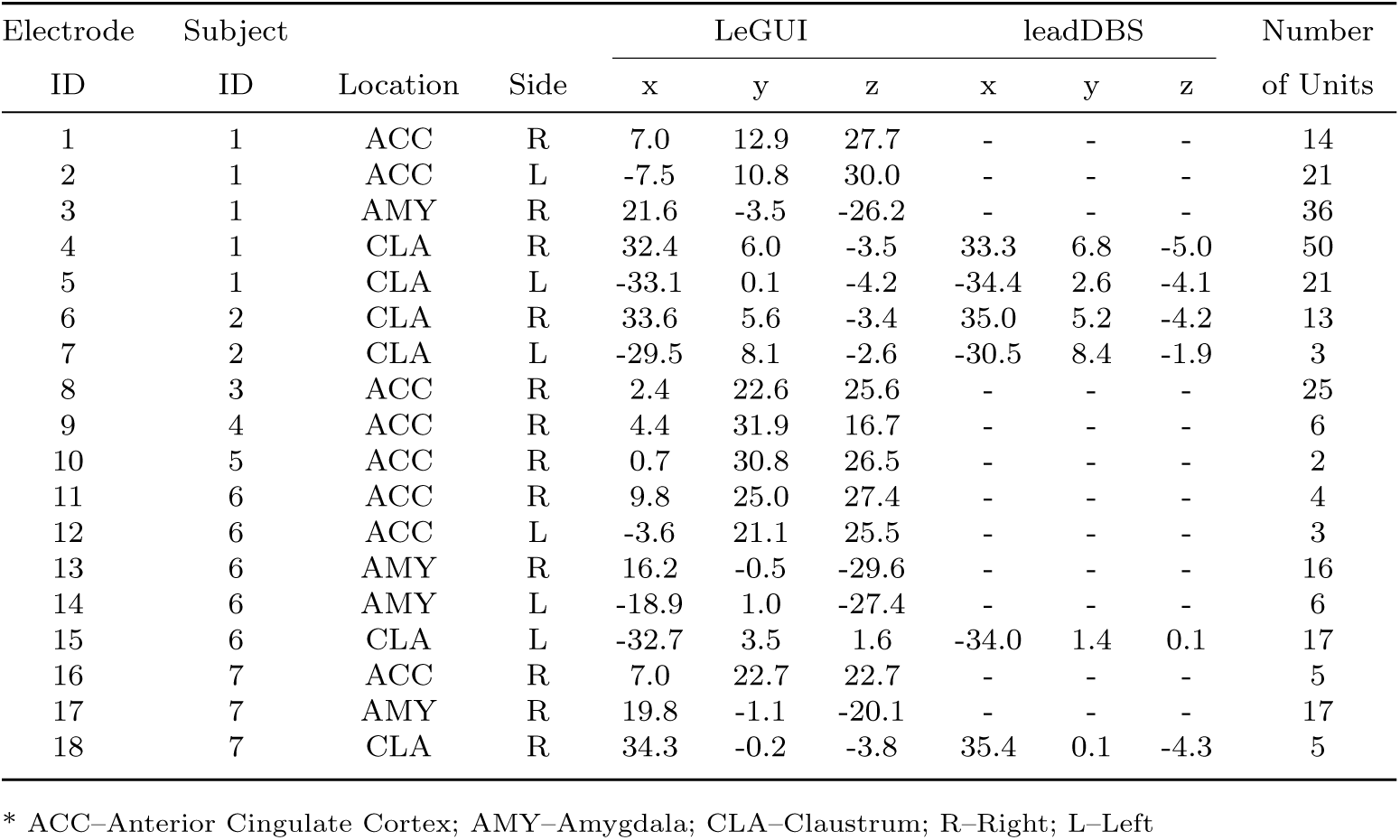
MNI coordinates of microwires for different brain regions.

| Electrode ID | Subject |  |  | LeGUI |  |  | leadDBS |  |  | Number of Units |
| --- | --- | --- | --- | --- | --- | --- | --- | --- | --- | --- |
|  | ID | Location | Side | x | y | z | x | y | z |  |
| 1 | 1 | ACC | R | 7.0 | 12.9 | 27.7 | - | - | - | 14 |
| 2 | 1 | ACC | L | -7.5 | 10.8 | 30.0 | - | - | - | 21 |
| 3 | 1 | AMY | R | 21.6 | -3.5 | -26.2 | - | - | - | 36 |
| 4 | 1 | CLA | R | 32.4 | 6.0 | -3.5 | 33.3 | 6.8 | -5.0 | 50 |
| 5 | 1 | CLA | L | -33.1 | 0.1 | -4.2 | -34.4 | 2.6 | -4.1 | 21 |
| 6 | 2 | CLA | R | 33.6 | 5.6 | -3.4 | 35.0 | 5.2 | -4.2 | 13 |
| 7 | 2 | CLA | L | -29.5 | 8.1 | -2.6 | -30.5 | 8.4 | -1.9 | 3 |
| 8 | 3 | ACC | R | 2.4 | 22.6 | 25.6 | - | - | - | 25 |
| 9 | 4 | ACC | R | 4.4 | 31.9 | 16.7 | - | - | - | 6 |
| 10 | 5 | ACC | R | 0.7 | 30.8 | 26.5 | - | - | - | 2 |
| 11 | 6 | ACC | R | 9.8 | 25.0 | 27.4 | - | - | - | 4 |
| 12 | 6 | ACC | L | -3.6 | 21.1 | 25.5 | - | - | - | 3 |
| 13 | 6 | AMY | R | 16.2 | -0.5 | -29.6 | - | - | - | 16 |
| 14 | 6 | AMY | L | -18.9 | 1.0 | -27.4 | - | - | - | 6 |
| 15 | 6 | CLA | L | -32.7 | 3.5 | 1.6 | -34.0 | 1.4 | 0.1 | 17 |
| 16 | 7 | ACC | R | 7.0 | 22.7 | 22.7 | - | - | - | 5 |
| 17 | 7 | AMY | R | 19.8 | -1.1 | -20.1 | - | - | - | 17 |
| 18 | 7 | CLA | R | 34.3 | -0.2 | -3.8 | 35.4 | 0.1 | -4.3 | 5 |

\* ACC–Anterior Cingulate Cortex; AMY–Amygdala; CLA–Caudate; R–Right; L–Left

**Table 3.**
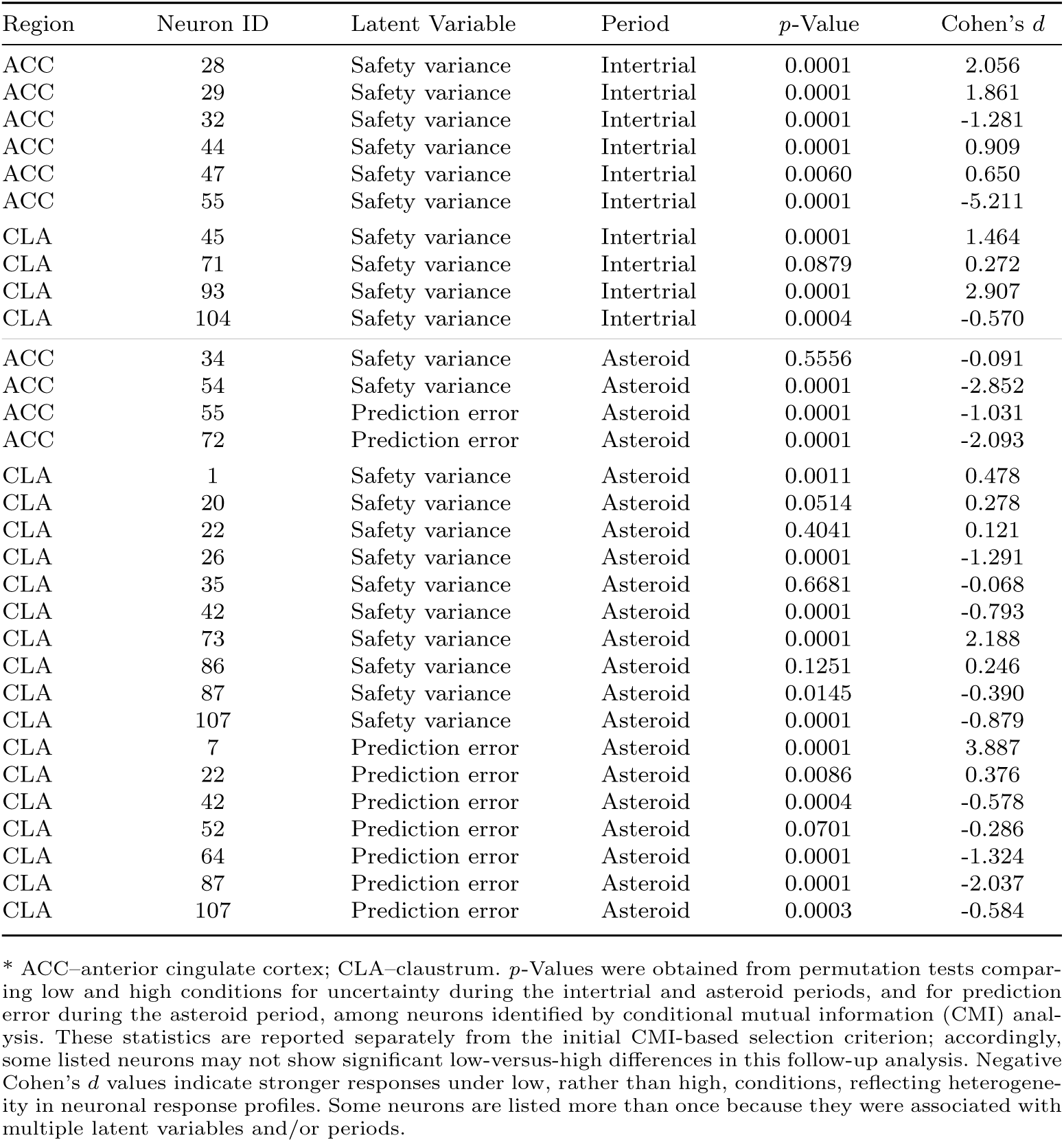
ACC and CLA *p*-values and Cohen’s *d* for low vs. high conditions for neurons identified by conditional mutual information analysis.

**Table 4.** Decoding performance by brain region, epoch, and latent variable.

| Brain Region | Epoch | Latent Variable | True: mean ± CI | $p$-value | Cohen's $d$ |
| --- | --- | --- | --- | --- | --- |
| ACC | intertrial | uncertainty | $0.508 \pm 0.011$ | 0.096 | 0.16 |
| ACC | intertrial | prediction error | $0.505 \pm 0.010$ | 0.119 | 0.14 |
| ACC | asteroid | uncertainty | $0.509 \pm 0.010$ | 0.032 | 0.22 |
| ACC | asteroid | prediction error | $0.512 \pm 0.010$ | 0.009 | 0.28 |
| AMY | intertrial | uncertainty | $0.506 \pm 0.008$ | 0.075 | 0.17 |
| AMY | intertrial | prediction error | $0.496 \pm 0.008$ | 0.824 | -0.11 |
| AMY | asteroid | uncertainty | $0.504 \pm 0.008$ | 0.128 | 0.13 |
| AMY | asteroid | prediction error | $0.499 \pm 0.008$ | 0.605 | -0.03 |
| CLA | intertrial | uncertainty | $0.505 \pm 0.007$ | 0.078 | 0.14 |
| CLA | intertrial | prediction error | $0.504 \pm 0.006$ | 0.103 | 0.12 |
| CLA | asteroid | uncertainty | $0.513 \pm 0.007$ | 0.0003 | 0.36 |
| CLA | asteroid | prediction error | $0.510 \pm 0.007$ | 0.004 | 0.25 |

\* ACC–Anterior Cingulate Cortex; AMY–Amygdala; CLA–Caudate; CI - 95% bootstrap CI (10,000 resamples)

**Table 5.**
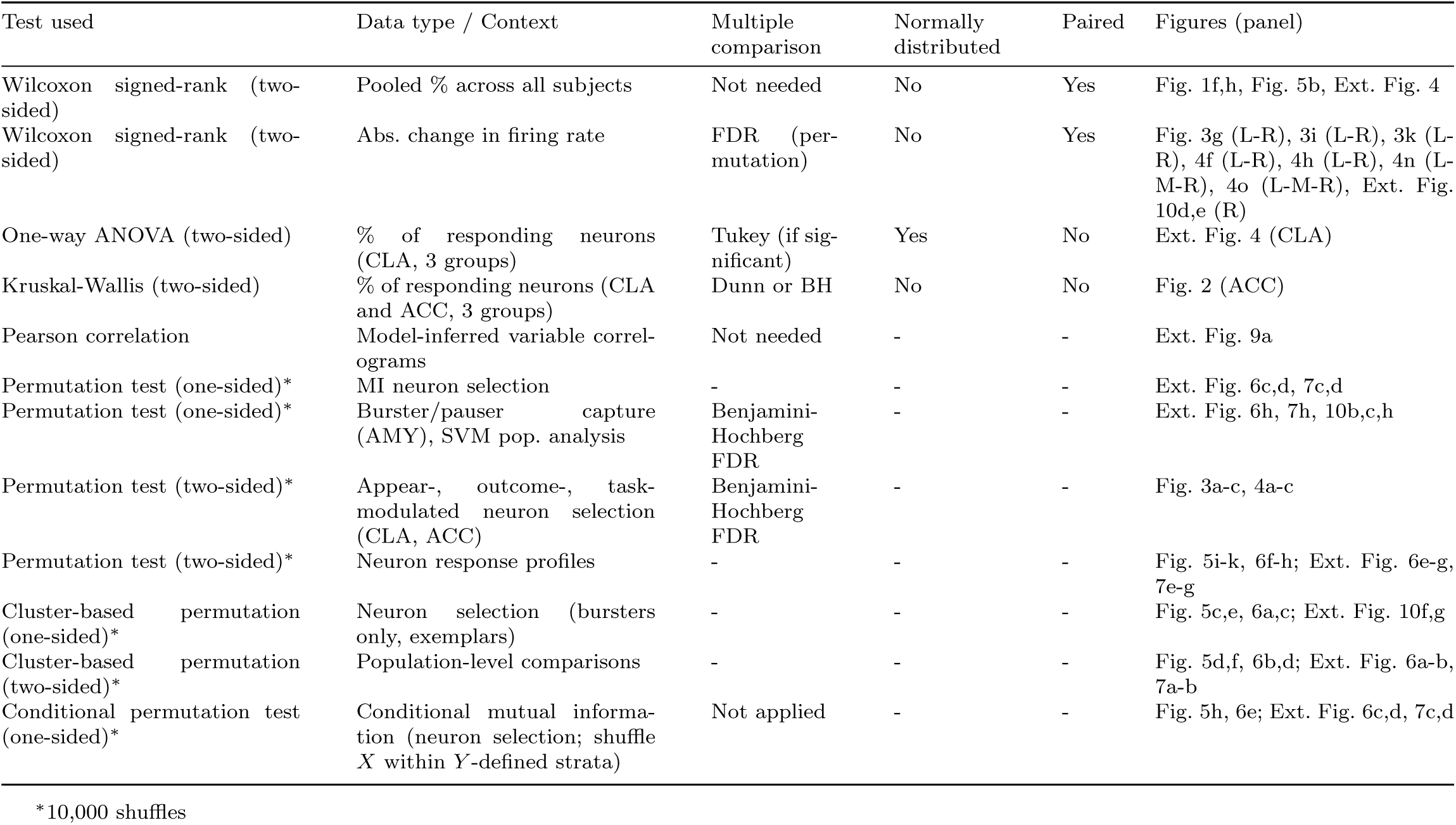
Summary of statistical tests.

**Extended Data Figure 1.**
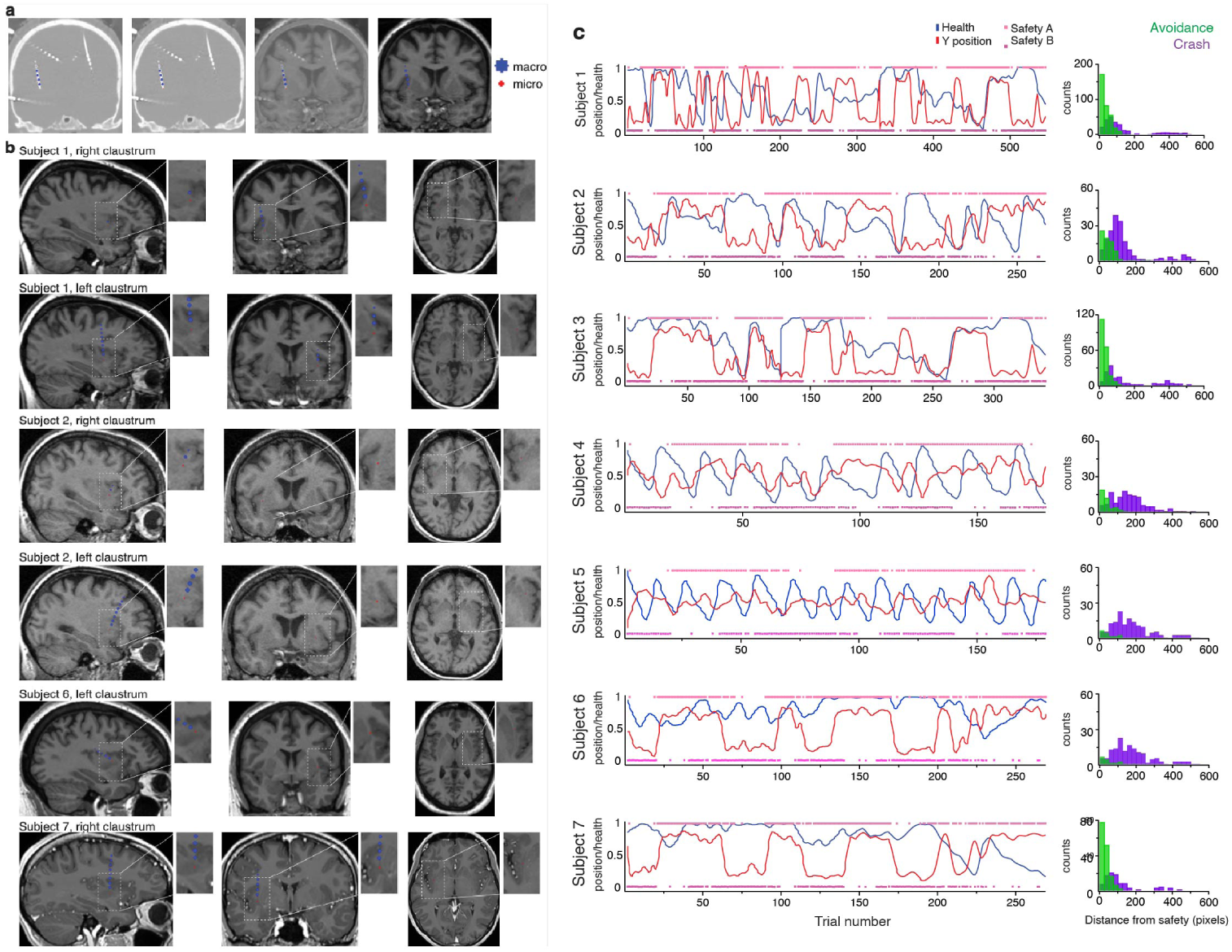
CLA Electrode location and task performance per patient. (a) Post-implantation CT and preoperative MRI scans showing the location of the microwire bundle (red cross) used to record CLA units, positioned at the tip of each macroelectrode (blue cross). From left to right: CT, two fused CT–MRI views, and MRI. (b) Subject-level fusion of post-implantation CT with preoperative MRI scans depicting the trajectory of macroelectrodes (blue cross) and the location of microelectrodes (red cross) within the claustrum for each patient. Subjects 3, 4, and 5 are not shown as no CLA electrode was implanted. (c) Task performance for all subjects. Left: subject-level traces showing spaceship *y*-position (red) and spaceship integrity (blue) throughout the task. Pink and magenta lines indicate the approximate locations of safety zones across trials (spatial jitter not shown). Right: distribution of spaceship *y*-position relative to the nearest safety zone at the time of asteroid appearance preceding crash (purple) or avoidance (green) outcomes, shown across all trials for each subject.

**Extended Data Figure 2.**
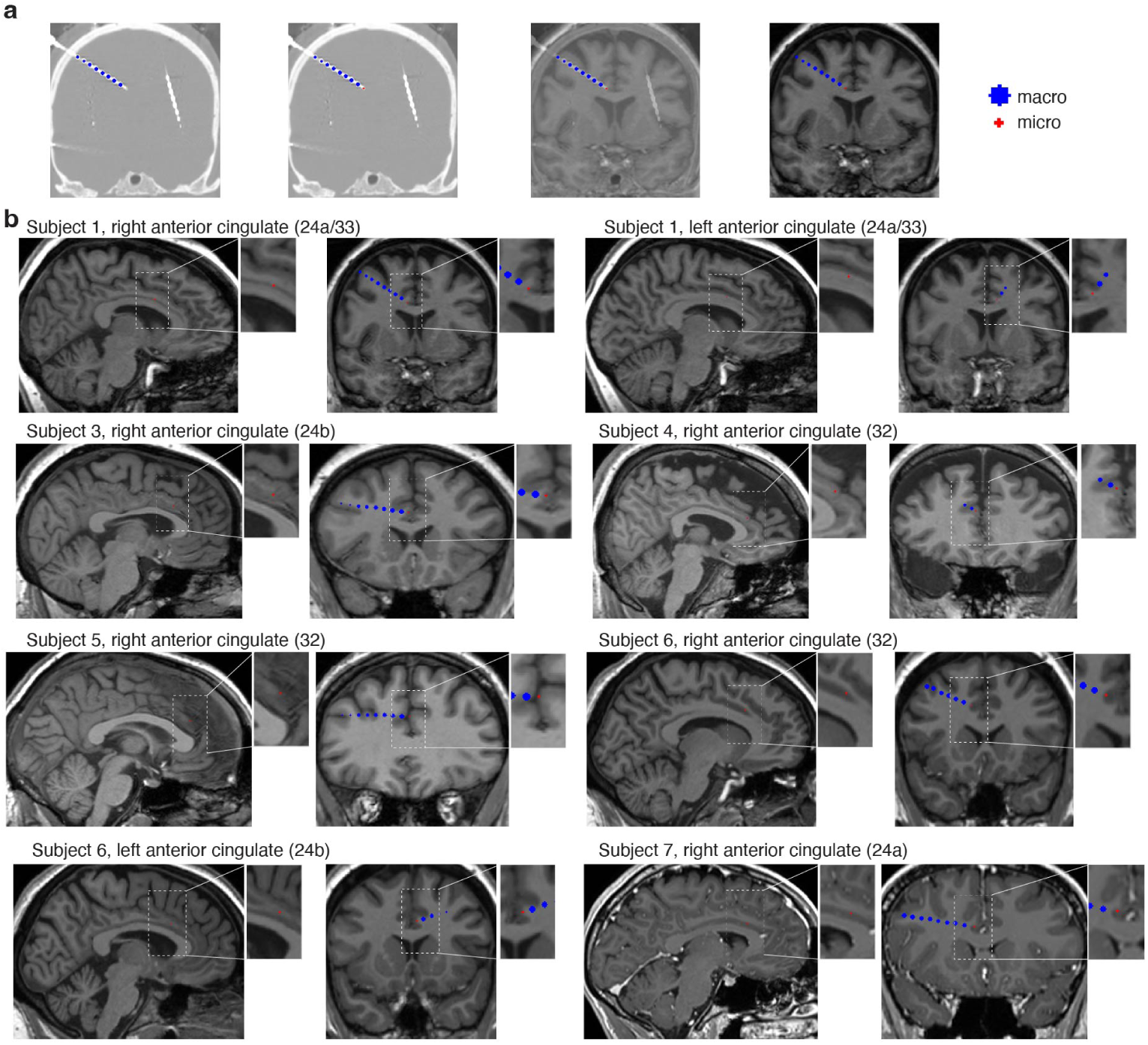
ACC Electrode location. (a) Post-implantation CT and preoperative MRI scans showing the location of the microwire bundle (red cross) used to record ACC units, positioned at the tip of each macroelectrode (blue cross). From left to right: CT, two fused CT–MRI views, and MRI (b) Subject-specific MRI views of ACC microwire localization. Each row displays two subjects, and for each subject two anatomical planes are shown (sagittal (left) and coronal (right)). Dashed boxes indicate the region of interest (ROI), with magnified insets highlighting the microwire position within the ACC. The targeted ACC subregion for each subject (areas: 24a/33, 24a, 32, and 24b; [78]) is indicated in the panel titles. Subject 2 is not shown as no ACC electrode was implanted.

**Extended Data Figure 3.**
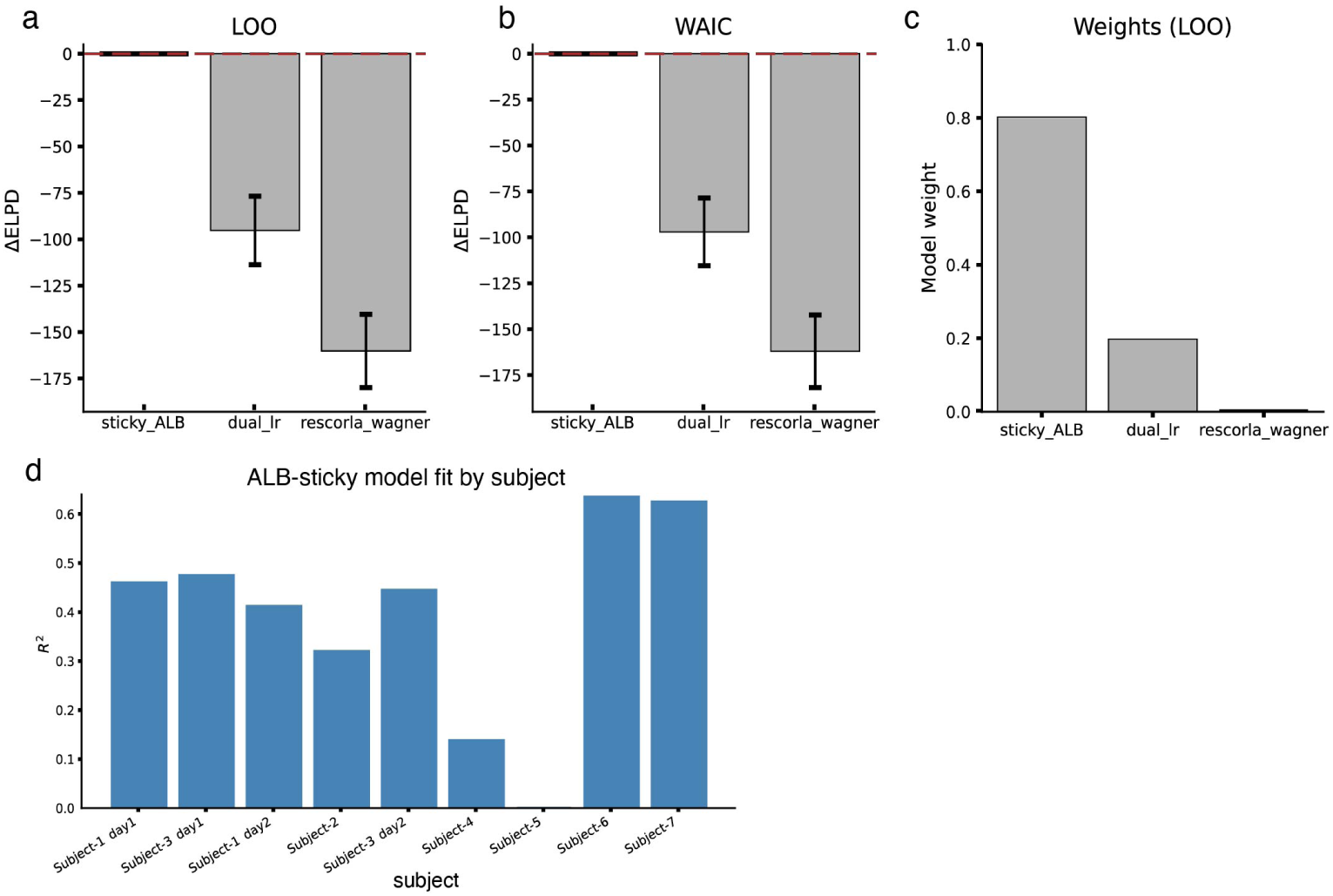
Model comparison and subject-level goodness-of-fit for ALB-sticky model of behavior in the spaceship task. (a,b) Comparison of candidate models using leave-one-out cross-validation (LOO; A) and the widely applicable information criterion (WAIC; B). Bars represent differences in expected log predictive density (ΔELPD) relative to the best-performing model, with error bars indicating the standard error of the difference. Values are shown relative to the best-performing model (red dashed line at 0), such that more negative values indicate worse predictive accuracy. Across both metrics, the ALB-sticky model showed superior predictive performance compared with the dual learning-rate and Rescorla–Wagner models. (c) LOO model weights likewise favored the ALB-sticky model. (d) Goodness-of-fit of the winning ALB-sticky model for individual subjects or sessions in the spaceship task, quantified using *R*^2^. These results support the use of the ALB-sticky model for deriving trial-wise latent variables in subsequent analyses.

**Extended Data Figure 4.**
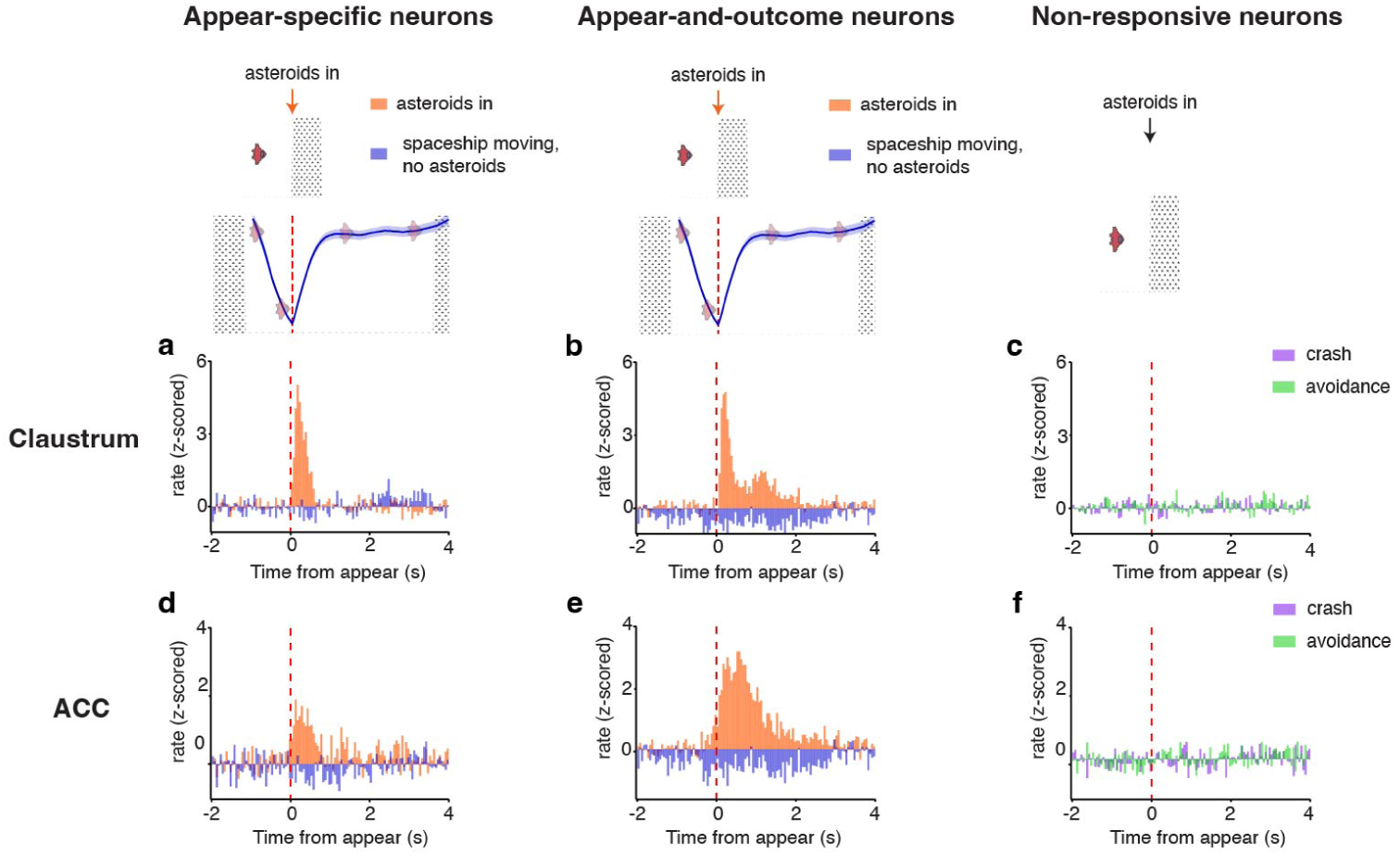
Appear-modulated CLA and ACC neurons during visual cues. (a) Peri-stimulus time histograms (PSTHs) of appearance-specific claustrum (CLA) neurons aligned to asteroid appearance (orange) versus spaceship movement in the absence of asteroids on the screen (blue). Appearance-specific neurons increased firing selectively during asteroid appearance, but not during spaceship movement (two-sided Wilcoxon signed-rank test comparing z-scored firing rates from 0 to 2 s post-event onset against zero; *n* = 23 neurons; asteroids: *Z* = 3.64, *p* = 2.62 *×* 10^−4^; spaceship movement: *Z* = *−*0.39, *p* = 0.69). (b) Same as (a), but for appearance– and outcome-modulated CLA neurons (two-sided Wilcoxon signed-rank test comparing z-scored firing rates from 0 to 2 s post-event onset against zero; *n* = 38 neurons; asteroids: *Z* = 1.80, (n.s.) *p* = 0.07; spaceship movement: *Z* = 0.97, *p* = 0.33). (c) PSTHs of non-responsive task neurons in the CLA aligned to asteroid appearance during crash (purple) or avoidance (green) trials. No significant responses were observed (two-sided Wilcoxon signed-rank test comparing z-scored firing rates from 0 to 2 s post-asteroid onset against zero; *n* = 32 neurons; crash: *Z* = 0.20, *p* = 0.84; avoidance: *Z* = 0.41, *p* = 0.68). (d) Same as (a), but for appearance-specific anterior cingulate cortex (ACC) neurons (two-sided Wilcoxon signed-rank test comparing z-scored firing rates from 0 to 2 s post-event onset against zero; *n* = 16 neurons; asteroids: *Z* = 2.48, *p* = 0.013; spaceship movement: *Z* = *−*1.55, *p* = 0.12). (e) Same as (b), but for ACC neurons (two-sided Wilcoxon signed-rank test comparing z-scored firing rates from 0 to 2 s post-event onset against zero; *n* = 27 neurons; asteroids: *Z* = 2.79, *p* = 0.005; spaceship movement: *Z* = *−*1.89, *p* = 0.058). (f) Same as (c), but for ACC neurons (two-sided Wilcoxon signed-rank test comparing z-scored firing rates from 0 to 2 s post-asteroid onset against zero; *n* = 24 neurons; crash: *Z* = *−*0.94, *p* = 0.35; avoidance: *Z* = *−*0.14, *p* = 0.88).

**Extended Data Figure 5.**
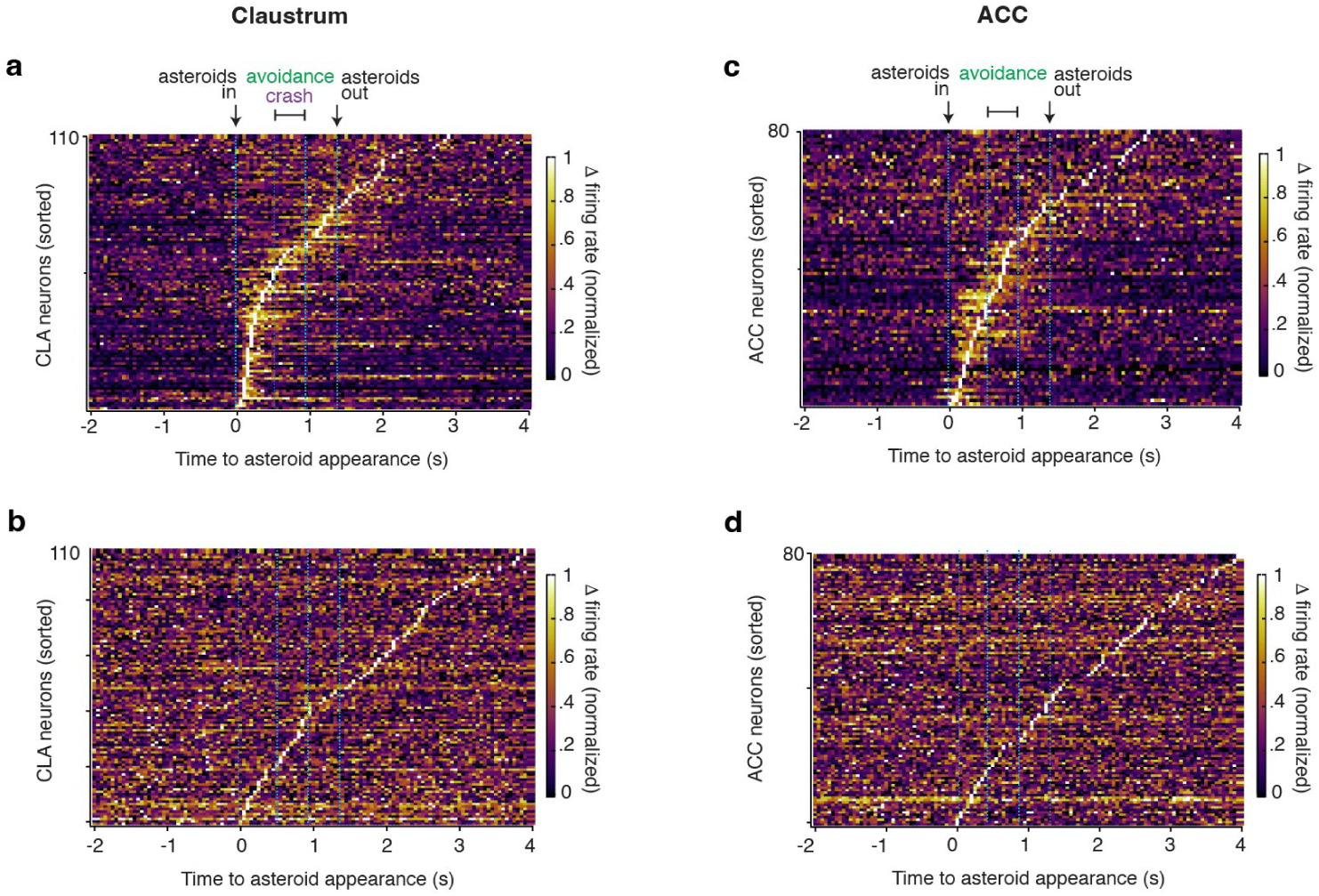
Sequence of activation of neurons after asteroid appearance in CLA and ACC. (a) Heatmap showing the normalized change in firing rate for all recorded claustrum (CLA) neurons (*n* = 110), aligned to asteroid appearance (0 s). Neurons are sorted by maximal firing rate change following asteroid appearance. Blue vertical dotted lines indicate the time intervals used for analysis. (b) Control heatmap displaying the same CLA neurons as in (a), but with temporally shuffled data. (c) Same as (a), but for all recorded anterior cingulate cortex (ACC) neurons (*n* = 80). (d) Same as (b), but for ACC neurons. Color scale indicates normalized change in firing rate, ranging from 0 (no change) to 1 (maximal change).

**Extended Data Figure 6.**
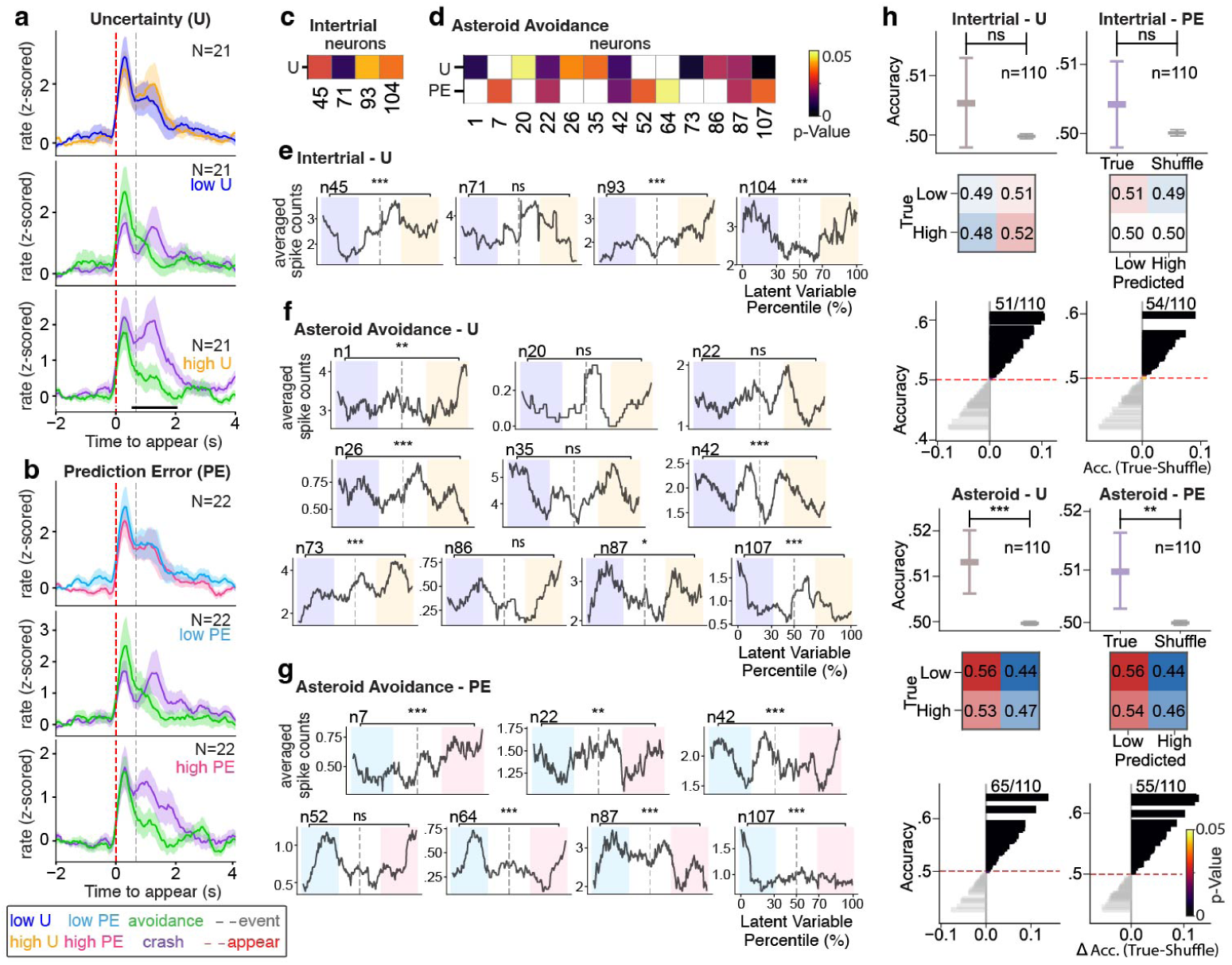
Extended analyses of uncertainty and prediction error in CLA neurons. (a–b) Z-scored firing rates (mean *±* s.e.m.) for neurons modulated by low uncertainty (a; *n* = 21) and low prediction error (b; *n* = 22), identified using the same selection criteria as the high uncertainty and high prediction error modulated neurons shown in Fig. 5d,f. Middle and bottom panels show responses for low and high uncertainty (a) and prediction error (b) trials, with each group further separated by action-contingent outcome (avoidance, green; crash, purple). In (a), bottom panel: t = 4.77. Black bars indicate time periods during which neuronal responses differed significantly between conditions (two-sided cluster-based permutation test, *p <* 0.05). (c–d) Heatmaps of neurons with significant conditional mutual information with latent variables during the intertrial (uncertainty) (c) and asteroid-avoidance (uncertainty and prediction error) (d) epochs. (e–g) Smoothed tuning curves were constructed by ranking trials according to the latent variable and plotting the corresponding trial-averaged spike counts as a function of this ranked value. Low and high conditions correspond to trials below the 30th percentile and above the 70th percentile, respectively. (e) Intertrial: uncertainty. (f) Asteroid-avoidance: uncertainty. (g) Asteroid-avoidance: prediction error. Statistical significance was assessed using two-sided permutation tests comparing mean firing rates in low versus high conditions (see Table S3). (h) Shows decoding results for each epoch (intertrial and asteroid avoidance) and each latent variable (uncertainty and prediction error). Top: Mean decoding accuracy for classifiers trained on true labels compared with label-shuffled controls (mean *±* 95% bootstrap CI; one-sided permutation test; *n* = 110 neurons; see Table S4). Middle: Row-normalized confusion matrices (true vs. predicted; Low/High; chance = 0.50). Bottom: Neuron-wise decoding performance showing accuracy (*y*-axis) and ΔAccuracy (True – Shuffle; *x*-axis); Gray bars denote neurons whose decoding performance did not exceed shuffled-label baseline, whereas neurons that exceeded shuffled controls are colored according to their FDR-corrected *p*-value colormap. Red dashed line indicates the nominal chance level (accuracy = 0.5).

**Extended Data Figure 7.**
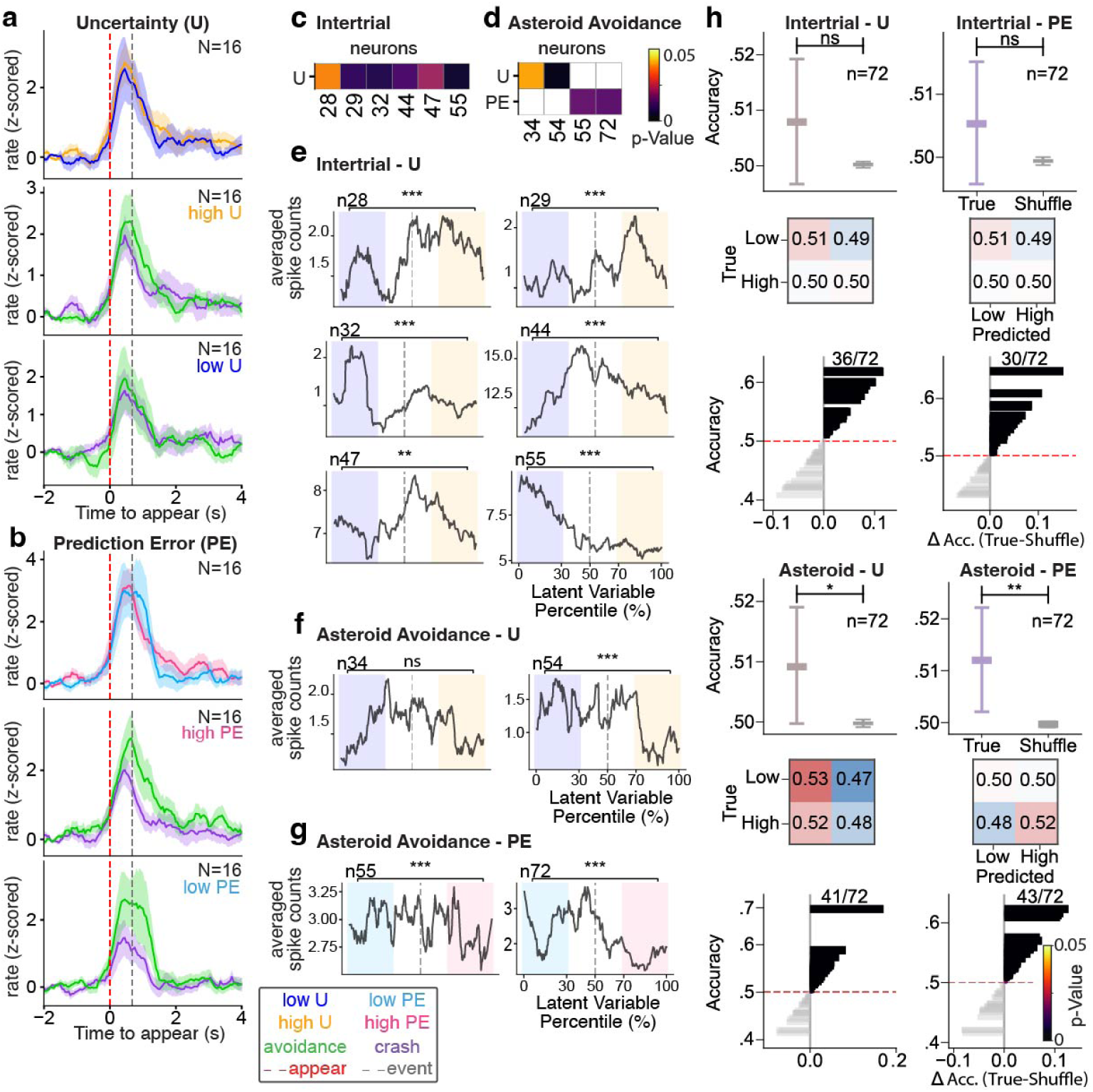
Extended analyses of uncertainty and prediction error in ACC. (a–b), Z-scored firing rates (mean *±* s.e.m.) for neurons modulated by high uncertainty (a, *n* = 16) and prediction error (b, *n* = 16), identified using the same selection criteria as the low uncertainty and low prediction error modulated neurons shown in Fig. 6b,d. Middle and bottom panels show neurons responsive to high and low uncertainty (a) and prediction error (b) trials grouped by outcome (avoidance: green; crash: purple). Black bars indicate time periods during which neuronal responses differed significantly between conditions, identified using two-sided cluster-based permutation test (*p <* 0.05). (c–d) Heatmaps of neurons with significant conditional mutual information during intertrial (c) and asteroid avoidance (d) periods. (e–g) Smoothed tuning curves were constructed by ranking trials according to the latent variable and plotting the corresponding trial-averaged spike counts as a function of this ranked value. Low and high conditions correspond to trials below the 30th percentile and above the 70th percentile, respectively. (e) Intertrial and (f) asteroid avoidance epoch uncertainty, and (g) asteroid avoidance epoch prediction error. Statistical significance assessed using two-sided permutation tests comparing mean firing rate in low vs high conditions (see Table S3). (h) Shows decoding results for each epoch (intertrial and asteroid avoidance) and each latent variable (uncertainty and prediction error). Top: Decoding accuracy for true labels and label-shuffled controls (mean *±* 95% bootstrap CI; one-sided permutation test; *n* = 72; see Table S4). Middle: Row-normalized confusion matrices (true vs. predicted; Low/High; chance = 0.50). Bottom: Neuron-wise decoding performance showing accuracy (*y*-axis) and ΔAccuracy (True – Shuffle; *x*-axis); Gray bars denote neurons whose decoding performance did not exceed shuffled-label baseline, whereas neurons that exceeded shuffled controls are colored according to their FDR-corrected *p*-value colormap. Red dashed line indicates the nominal chance level (accuracy = 0.5).

**Extended Data Figure 8.**
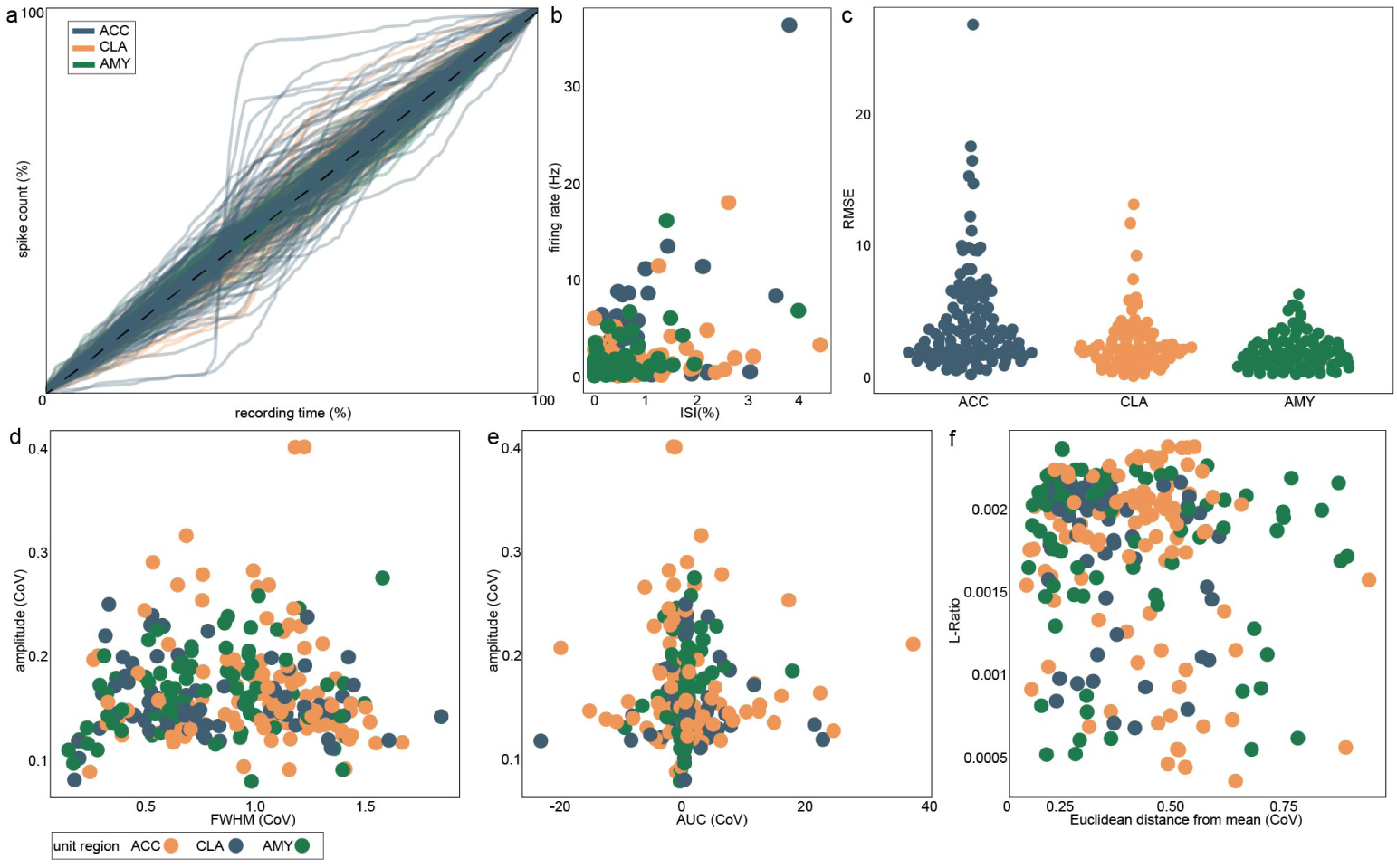
Metrics of single-unit stability. (a) Cumulative distribution of total detected spikes (%) over the recording session (%), colored by unit region. (b) Relationship between firing rate (Hz) and inter-spike interval (ISI) violations (% of spikes with inter-spike intervals *<* 3 ms), with points colored by brain region (ACC, CLA, or AMY). (c) Dot plot of the root mean squared error (RMSE) of single-unit cumulative distributions relative to the dashed diagonal reference line in panel (a), stratified by unit region. (d–f) Scatterplots of waveform-based stability metrics computed from randomly sampled spikes. (d) Coefficient of variation (CoV) of maximal waveform amplitude versus CoV of full width at half maximum (FWHM). (e) CoV of maximal waveform amplitude versus CoV of area under the curve (AUC). (f) L-ratio versus CoV of Euclidean distance from the mean waveform. All points are colored by recording region (ACC, CLA, or AMY).

**Extended Data Figure 9.**
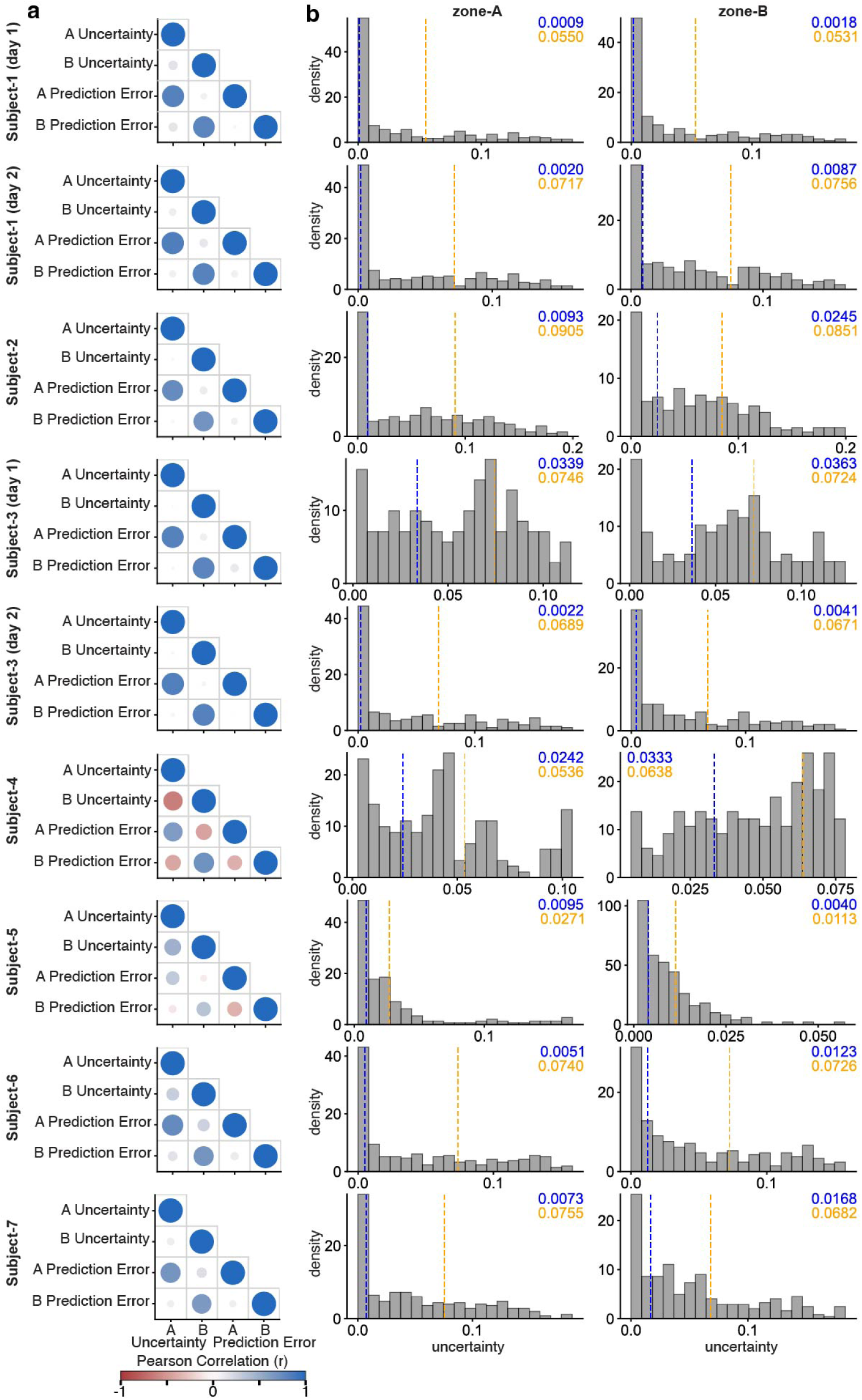
Subject-wise model-inferred variable correlograms and metrics of model-inferred uncertainty. (a) Each panel displays the pairwise Pearson correlation coefficients between A and B uncertainty and prediction error measures (associated with safety zones A and B). The color of each circle indicates the sign and strength of the correlation, and the size of each circle reflects the absolute correlation magnitude. Behavioral labels are shown along the axes. Most subjects showed predominantly positive correlations, whereas Subject 4 exhibited some negative correlations, particularly among prediction error measures. (b) Data from individual subjects. Probability-density histograms (20 bins) of trial-wise subjective uncertainty for each subject. Vertical dashed lines indicate the thresholds used to define high (orange; 70th percentile) and low (blue; 30th percentile) uncertainty conditions. For Subjects 1 and 3, days 1 and 2 are shown separately.

**Extended Data Figure 10.**
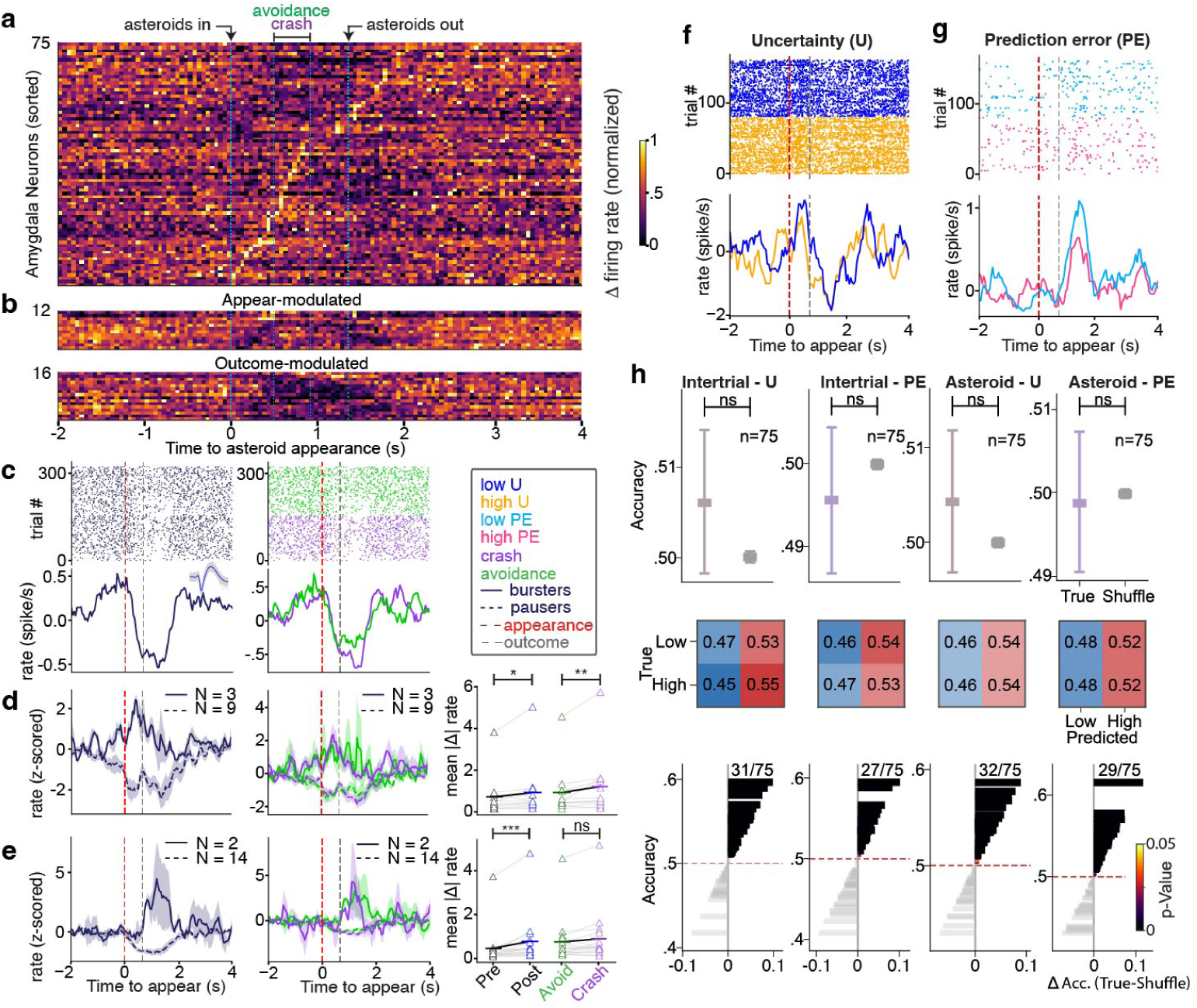
Amygdala neuronal response to task and modulation by uncertainty and prediction error. (a) Heatmap of normalized firing rates for all amygdala (AMY) neurons (*n* = 75) sorted by latency to maximal firing-rate change (blue dotted line, time used for analysis). (b) Heatmaps of appearance-modulated AMY neurons (top; *n* = 12; 3 bursters, 9 pausers) and outcome-modulated AMY neurons (bottom; *n* = 16; 2 bursters, 14 pausers). (c) Example of appear– and outcome-modulated AMY neuron. Spike rasters (top) and mean firing rate (bottom) for all trials (left) and by outcome (right; avoidance, green; crash, purple). (d–e) Left panel, average response (z-score *±* s.e.m.) for appear– (d) and outcome-modulated (e) AMY neurons. Middle panel, average response (z-score *±* s.e.m.) split by outcome (avoidance vs. crash). Bursters (solid line: d, *n* = 3; e, *n* = 2) and pausers (dashed line: d, *n* = 9; e, *n* = 14) are shown. Right panel, mean absolute firing-rate change; left, pre– vs. post-appearance (d, *n* = 12, *Z* = 2.04, *p* = 0.0425; e, *n* = 16, *Z* = 3.52, *p* = 0.0001) and right, avoidance vs. crash (d, *n* = 12, *Z* = 2.59, *p* = 0.007; e, *n* = 16, *Z* = 2.02, *p* = 0.05). (f–g) Examples of AMY neurons modulated by uncertainty (f: high, orange; low, blue) and prediction error (g: high, pink; low, light blue). For (f–g), spike rasters (top) and firing rate (bottom) are shown. (h) Columns show decoding results for latent variable (uncertainty and prediction error) within epoch (intertrial and asteroid). Top: Mean decoding accuracy for true labels and label-shuffled controls (mean *±* 95% bootstrap CI; one-sided permutation test; *n* = 75; see Table S4). Middle: Row-normalized confusion matrices (true vs. predicted labels), with the color scale centered at chance performance (0.5). Bottom: Neuron-wise decoding performance showing decoding accuracy (*y*-axis) versus improvement over shuffled control (ΔAccuracy = True – Shuffle; *x*-axis; FDR-corrected); Gray bars denote neurons whose decoding performance did not exceed shuffled label baseline; neurons that exceeded shuffled controls are colored by a p-value colormap. The dashed line indicates chance accuracy.

